# Decoding sexual dimorphism of the sex-shared nervous system at single-neuron resolution

**DOI:** 10.1101/2024.12.27.630541

**Authors:** Rizwanul Haque, Hagar Setty, Ramiro Lorenzo, Gil Stelzer, Ron Rotkopf, Yehuda Salzberg, Gal Goldman, Sandeep Kumar, Shiraz Nir Halber, Andrew M. Leifer, Elad Schneidman, Patrick Laurent, Meital Oren-Suissa

## Abstract

Sex-specific behaviors are often attributed to differences in neuronal wiring and molecular composition, yet how genetic sex shapes the molecular architecture of the nervous system at the individual neuron level remains unclear. Here, we use single-cell RNA sequencing to profile the transcriptome of sex-shared neurons in adult *Caenorhabditis elegans* males and hermaphrodites. We uncover widespread molecular dimorphism across the nervous system, including in previously unrecognized neuron-types such as the touch receptors. Neuropeptides and signaling-related genes exhibit strong sex-biased expression, particularly in males, reinforcing the notion that neuropeptides are crucial for diversifying connectome outputs. Despite these differences, neurotransmitter identities remain largely conserved, indicating that functional dimorphism arises through modulatory, not identity-defining, changes.

We show that sex-biased expression of neurotransmitter-related genes correlates with bias in outgoing synaptic connectivity and identify regulatory candidates for synaptic wiring, including both shared and sex-specific genes. This dataset provides a molecular framework for understanding how subtle regulatory differences tune conserved circuits to drive sex-specific behaviors.

## Introduction

The molecular mechanisms that underlie the development of sex differences in the nervous system have been the focus of many studies in the past decade, especially due to the sex bias demonstrated in the prevalence and progression of several neurological diseases, such as anxiety and Parkinson’s disease(*1–3*). Recent studies have made an effort to dissect sexually dimorphic neuronal circuits, uncovering unique circuits across the animal kingdom that operate differently between the sexes to drive behavior (termed “sexually dimorphic”); such as the circuit for empathic behaviors in mice, aggressive behaviors in fruit flies and nociceptive and mechanosensory behaviors in worms(*4–7*). While these studies have provided valuable insights into how specific circuits are differentially shaped, a deeper understanding of the molecular repertoire of the nervous system in both sexes is still missing. With the advancement of RNA sequencing, molecular profiles of both sexes in mice were generated, discovering differentially expressed genes between the sexes in specific neuronal populations such as the medial amygdala (MeA) and the ventromedial nucleus of the hypothalamus (VMH)(*8–11*). Additional studies performed transcriptomic profiling of different brain regions while delineating the effect of circulating hormones on sex differences in gene expression(*12–14*). Although these studies provided insights regarding sexually dimorphic behaviors, they did not explore the contribution of specific genes to these behaviors. Additionally, the hormonal regulation in mammalian systems might mask the direct effect of sexual identity (the chromosome complement) on distinct molecular signatures at the level of single neurons.

In the nematode *Caenorhabditis elegans* (*C. elegans*), the genetic sex of each somatic cell, including neurons, is determined in a cell-autonomous manner: the ratio of sex-chromosomes to autosomes controls a transcriptional cascade which results in the expression of either hermaphrodite-specific or male-specific genes(*15*, *16*). Recent studies explored the contribution of the genetic sex to nervous system development, with insights into synaptic connectivity, circuitry, function, and behavior(*7*, *17–19*). Despite all these efforts, it remains unclear to what extent does the genetic sex shapes gene expression patterns across the nervous system and the different neuronal classes.

Recently, single-cell RNA sequencing (scRNA-seq) approaches resolved the molecular heterogeneity of the hermaphrodite nervous system. Its 302 neurons were subdivided into 128 transcriptionally distinct neuron types(*20*). The transcriptomic signatures of each neuron class facilitated the identification of markers, unraveled mechanisms of cell specification, and enabled deeper characterization of nervous system development and aging(*20–29*). Earlier scRNA-seq focused only on hermaphrodites (with one exception for glia(*30*)), limiting our knowledge of the molecular signatures of male neurons and, most importantly, the differences in molecular signatures between the sexes. To gain a broad perspective on the molecular landscape of the nervous system in both sexes of *C. elegans*, we generated comparative scRNA-seq transcriptomic profiles for the sex-shared neuron types in both males and hermaphrodites (https://www.weizmann.ac.il/dimorgena/).

Our research provides an atlas for expression patterns of 18,975 genes in the nervous system of adult males and hermaphrodites. This atlas revealed a variety of neuronal gene classes with unique dimorphic gene expression patterns, with most of the neuropeptide-encoding genes demonstrating robust sexually dimorphic gene expression. We focused on the core, sex-shared nervous system i.e., neurons that exist in both sexes, and uncovered several sex-shared neurons with unique dimorphic transcriptomic profiles, including the sensory touch receptor neuron PLM. We harnessed our dataset to predict genes that might regulate connectivity and revealed gene candidates that operate in both sexes and in a sex-specific manner to control synaptic wiring. We further show that while neurotransmitter and terminal selector identity are broadly preserved, sexually dimorphic expression of neuromodulatory genes correlates with differences in neuronal output.

## Results

### Single-cell sequencing of adult hermaphrodite and male nervous system

To generate a single-cell transcriptomic profile of the nervous system in both sexes of *C. elegans* we dissociated young adult males and hermaphrodites expressing TagRFP pan-neuronally into single cells (fig. S1A). To generate pure male populations (XO animals), we used our protocol that enables large-scale isolation of males with high purity(*31*). To isolate the fluorescently labeled neurons, we dissociated young adult animals into single cells and sorted the single-cell suspensions by Fluorescence-Activated-Cell-Sorting (FACS) (Fig. 1A, fig. S1B-C, *See Methods*). We confirmed the neuronal enrichment and the intact morphology of RFP+ FACS-purified cells (fig. S1D-G, *See Methods*). Single-cell RNA sequencing was done using a microfluidic-based 10x genomics platform. We corrected for ambient RNA, removed low-quality and damaged cells, and performed cell-quality filtering (*See Methods*). This yielded a median of 925 unique molecular identifiers (UMIs) and 358 genes per cell after filtering. To filter neurons from non-neuronal cells, we used a cutoff based on known molecular landmarks (563 neuronal and 114 non-neuronal genes, table S1). After the removal of cells containing more than 20% of UMIs originating from mitochondrial genes, we obtained 13015 single-cell neuronal transcriptomes for hermaphrodites and 19454 single-cell transcriptomes for males, taken for further analysis and cluster annotation. To simplify cluster annotation, we merged our datasets with the previously annotated CeNGEN dataset(*20*). This combined dataset was first segregated into 72 partitions (fig. S2A). Sub-clustering of partition 1 (41219 cells) and partitions 2-72 (61546 cells) yielded 118 clusters (fig. S2B-C). A further iteration included manual assignment of varying identities, producing an overall of 156 clusters (details in *Methods*). 16 clusters were only present in males and 13 were classified as “unknown ID” due to lack of previous data. We annotated clusters using previously assigned cell identities(*20*) and manually assigned neuronal identities to 111 sex-shared clusters that correspond to 109 neuronal classes out of 116 (table S2, Fig. 1B)(*32*). We didn’t use male clusters, for which hermaphrodite clusters were not well represented in our data. 83 sex-shared clusters had more than 5 cells per cluster in both sexes (corresponding to 84 neuronal classes), leaving 28 sex-shared clusters that are underrepresented in at least one sex (low sample sizes), such as PHA and PHB (See table S2 for the full list). In total, in males, 93 sex-shared clusters contain more than 5 cells, including ASER, ASEL, AWC-OFF, and AWC-ON. The UMAP representations of sex-shared clusters of hermaphrodites (adult and L4(*20*)) and adult males showed a large overlap (Fig. 1C). When comparing UMAPs of hermaphrodites (YA vs L4), we observed a subtle shift, likely corresponding to the difference in their developmental stage (Fig. 1C, fig. S3)(*31*).

**Fig. 1:**
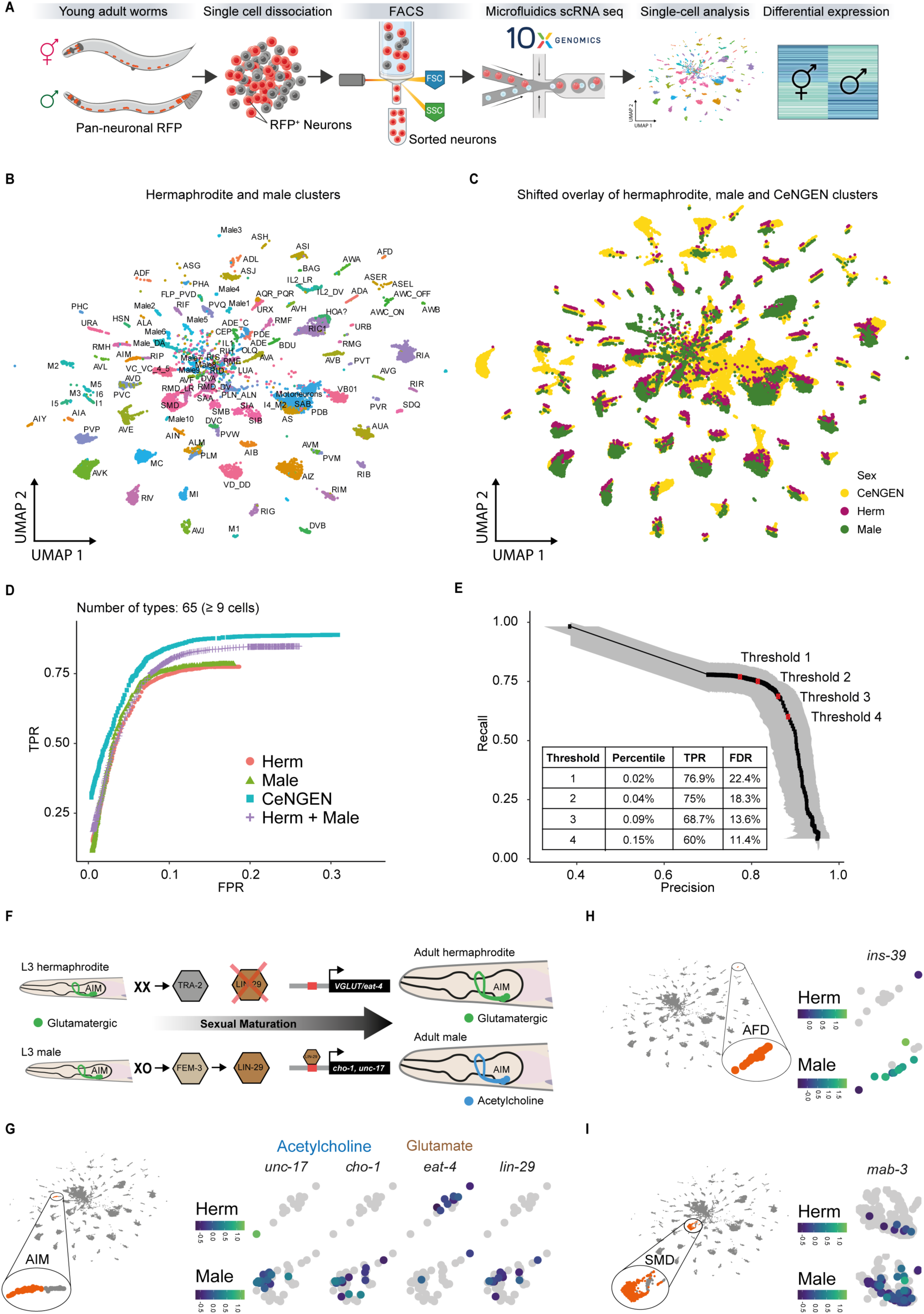
Single-cell RNA sequencing in both sexes of *C. elegans*. (A) Schematic of the experimental design. Hermaphrodite and male *C. elegans* harboring the pan-neuronal marker *rab-3p(prom1)::2xNLS::TagRFP* were subjected to cell dissociation. RFP+ cells were purified by FACS. The single-cell RNA transcriptomes were obtained using 10X Genomics (https://www.10xgenomics.com). (B) UMAP representation of all cell-clusters identified with their neuronal identity. Previously published(*20*) (CeNGEN) datasets or manual assignments were both used to determine the IDs of the neurons. (C) UMAP representation of cell clusters with a shifted overlay to illustrate the overlap. Data sets are YA hermaphrodites (magenta), YA males (green), and CeNGEN L4 hermaphrodites (yellow). (D) Receiver-Operator Characteristic (ROC) curve showing True Positive Rate (TPR) versus False Positive Rate (FPR) for hermaphrodites, males, hermaphrodites-males together, and CeNGEN compared to ground truth expression data using the 65 cell types with 9 or more cells in hermaphrodite and male clusters. (E) Precision-Recall (PR) Curve of Recall (TPR) versus Precision [1 – False Discovery Rate (FDR)] curve. Red dots represent thresholds 1-4. Grey shading represents 95% confidence intervals. (F) Schematic of AIM neurons neurotransmitter switch from glutamatergic identity (in hermaphrodite) to cholinergic identity (in male). (G) AIM representation in the UMAP projection of all cells (inset marked as orange). UMAP projection depicting expression of *unc-17*, *cho-1* cholinergic identity genes, *eat-4* glutamatergic identity gene, and transcriptional factor *lin-29* gene within AIM cluster. (H) High expression of *ins-39* in males. AFD representation in the UMAP projection of all cells (inset marked in orange). UMAP projection depicting expression of *ins-39* gene within AFD cluster. (I) Dimorphic expression of *mab-3*. Left, SMD representation in the UMAP projection of all cells (inset marked in orange). UMAP projection depicting expression of *mab-3* gene within SMD cluster. In G-I, heatmaps represent expression levels (log10). Created in BioRender. Oren, M. (2025) https://BioRender.com/i2zd1il.

### Single-cell profiles of the nervous system capture dimorphic gene expression

In this work, we focused on the sex-shared clusters and did not further cluster the male-specific neurons (table S3). For gene expression analysis, we calculated the fraction of cells expressing a gene in each cluster, the average expression, and markers by using Seurat(*33*) with default parameters (*See Methods*, https://www.weizmann.ac.il/dimorgena/). We also calculated expression thresholds as previously described ((*20*), *See Methods*) and combined this information with the cell type-marker analysis. Expression patterns across all neurons in adult males and hermaphrodites can be mined using one of four thresholds of increased stringency (Fig. 1D-E, table S3). We characterized the expression of genes with a known sex-biased expression. The Zn finger transcription factor LIN-29 promotes a fate switch from a glutamatergic (hermaphrodite) to a cholinergic (male) identity in the AIM neuron(*34*, *35*) (Fig. 1F). Indeed, AIM cell cluster showed expression of *lin-29*, the Choline Acetyltransferase *cho-1*/CHOT, and the vesicular Ach transporter *unc-17/VACHT* in male clusters, while hermaphrodite cluster expressed the vesicular glutamate transporter *eat-4/VGLUT* (Fig. 1G). We also observed sex-biased expression of *ins-39* in male AFD neurons and of *mab-3* in male SMD neurons, as previously reported (Fig. 1H-I)(*31*, *34*, *36*).

### Comparative analysis of sex-shared neuronal clusters reveals extensive and diverse molecular dimorphism

To uncover to what extent the transcriptomic profiles of the sex-shared nervous system diverge between males and hermaphrodites, we took a comparative approach, ranking the sex-shared neuronal clusters according to the number of differentially expressed genes (DEGs) (*See Methods*). For ranking purposes, we filtered out clusters with more than one identity, used a cut-off of 9 cells or more per cluster in both sexes (fig. 2SD), and proceeded with 62 sex-shared clusters (*See Methods*). This analysis captured known sex-shared neurons with dimorphic gene expression profile, including PHC, AFD, AIM, and AVG(*17*, *34–37*) (Fig. 2A). Strikingly, it also revealed many sex-shared neurons that were not previously classified as dimorphic, such as the six touch-receptor-neurons (TRNs), BDU, PVR interneurons and many more (Fig. 2A). Notably, all 62 analyzed sex-shared clusters contained a subset of significant DEGs, highlighting the widespread presence of dimorphic gene expression profile across the worm’s nervous system (Fig. 2A).

**Fig. 2.**
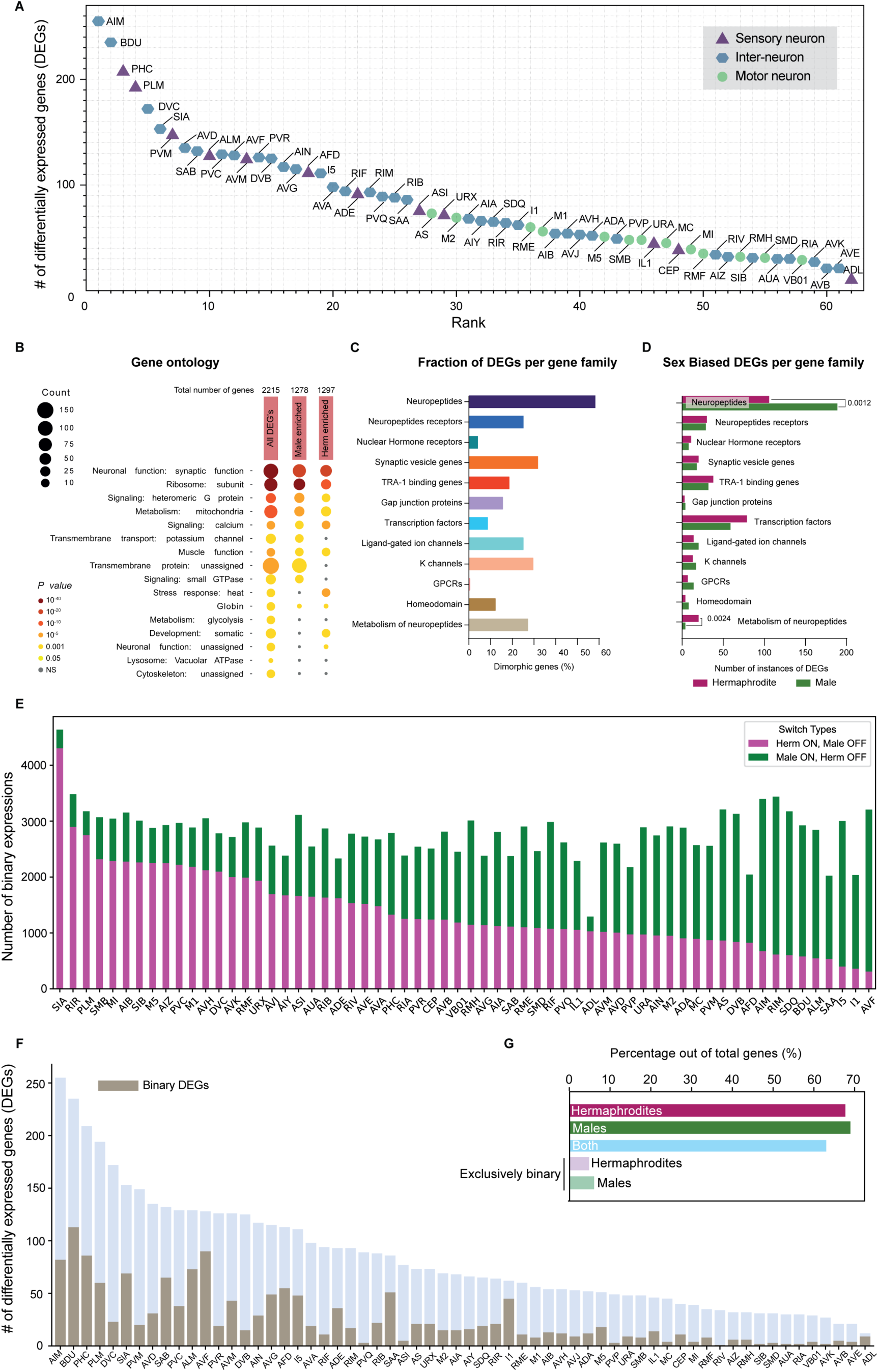
Comparative analysis of sex-shared neurons reveals widespread molecular dimorphism. (A) Number of differentially expressed genes (DEGs) in the sex-shared neuronal clusters. Clusters are divided to sensory (triangle), interneuron (hexagon), and motor (circle) neurons. A gene was considered DE with average log fold change > 1, ≥ 9 cells per cluster in both sexes, p-value < 0.05, and a threshold ≥ 2 in at least one sex (*See Methods*). (B) Gene ontology (GO) term analysis for the 2215 DEGs in both sexes and in hermaphrodite and males separately (only genes meeting the criteria of log fold change > 1, ≥ 9 cells per cluster in both sexes, p-value < 0.05, threshold ≥ 2 in at least one of the sexes are included) reveals specific enrichment for synaptic and signaling gene expression. Wormcat p-values are determined by one-sided Fisher test with FDR correction. (C) Percentage of DEGs from each gene family as a fraction of the number of genes in the specific family. (D) The number of instances of DEGs in the gene families presented in (C) divided according to DEGs that are enriched in males (green) and hermaphrodites (magenta). Only significant comparisons are shown. A DEG that is observed in more than one cluster was counted accordingly (*See Methods*). For (D) we performed a two-sided Fisher’s exact test followed by p-value Bonferroni correction for multiple comparisons. (E) Number of binary genes (‘on’ in hermaphrodites ‘off’ in males in magenta, ‘on’ in males ‘off’ in hermaphrodites in green) in each cell. Cells are shown according to the number of hermaphrodite binary genes from left to right. (F) Number of DEGs across cells (light blue), with the number of binary genes (gray). (G) Percentage of binary genes in hermaphrodites, males, both, exclusively (across cell types) hermaphrodites and exclusively males.

The ranking analysis identified a total of 2215 DEGs across all sex-shared clusters. Gene ontology (GO) enrichment analysis for these DEGs revealed a significant proportion of enriched terms related to neuronal function, suggesting their potential relevance to neuronal activities (Fig. 2B). Additional GO terms such as synaptic function, heteromeric G protein, and calcium signaling underscore the representation of genes that may play a role in shaping a dimorphic nervous system (Fig. 2B). By exploring neuronal DEGs based on their gene family(*38–43*), we found that while multiple gene families differ between the sexes, neuropeptides, neuropeptide receptors, and neuropeptide metabolism-related genes are overrepresented with a significant male-biased expression of neuropeptides and neuropeptides metabolism-related genes (58.5%, 92 of the 157 predicted neuropeptide genes are DEGs) (Fig. 2C-D, Fig. 3A, fig. S4A-B).

**Fig. 3.**
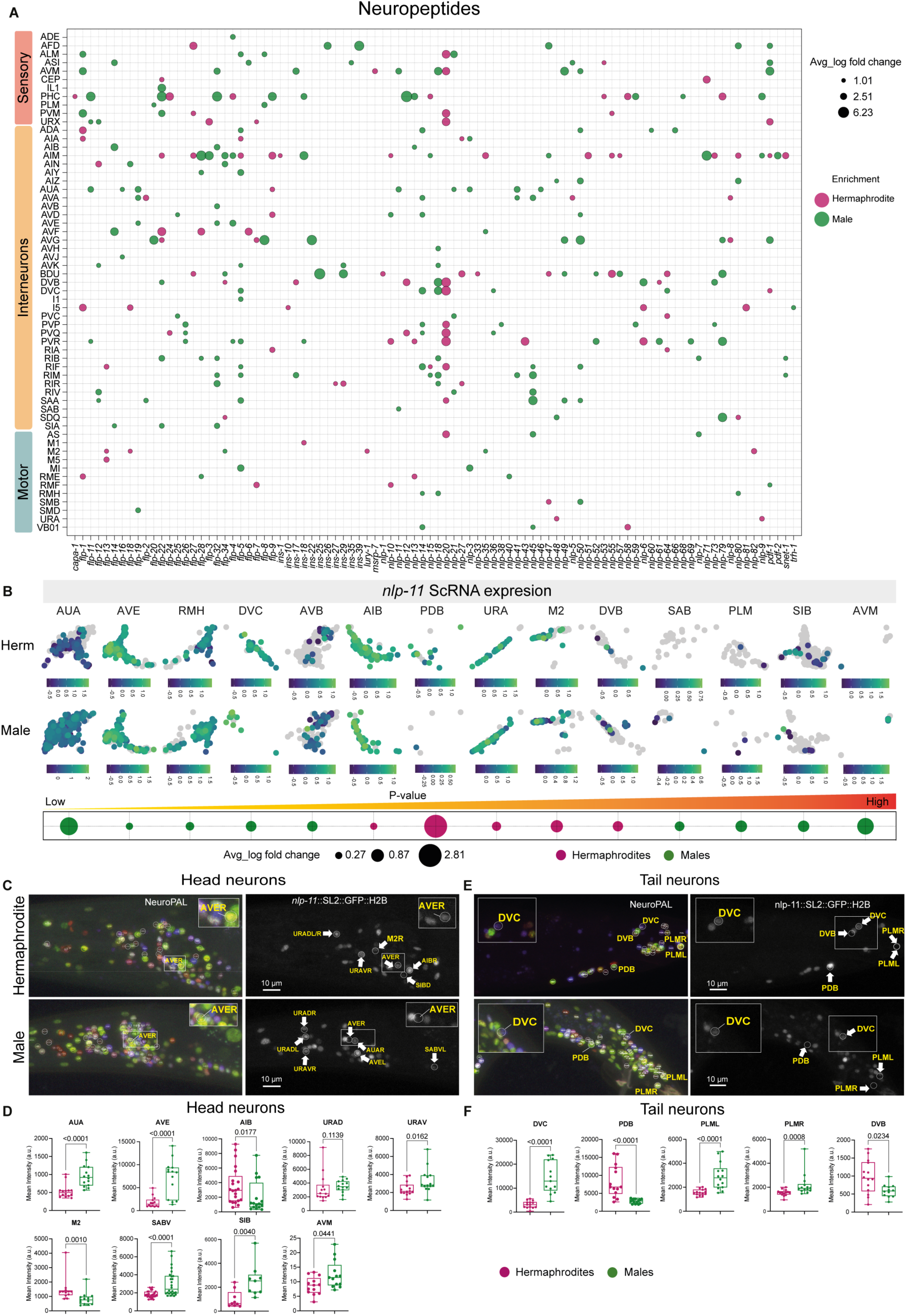
Sexual dimorphism of neuropeptides across the nervous system. (A) Bubble plot of differentially expressed neuropeptide genes between sexes. Bubble size indicates average fold change in gene expression (only genes with log fold-change >1, ≥9 cells/cluster, p <0.05, threshold ≥2 in at least one sex included), and bubble color shows sex enrichment (green: male, magenta: hermaphrodite). Rows represent neuron types (sensory, interneurons, and motor). Columns represent genes. (B) UMAP projection of AUA, AVE, RMH, DVC, AVB, AIB, PDB, URA, M2, DVB, SAB, PLM, SIB and AVM clusters showing expression of *nlp-11* gene in both sexes. Heatmap shows expression levels. Bubble plot representation of *nlp-11* gene expression in clusters as in (A) of two sexes, arranged left to right by p-value (most to least significant). Bubble size represents average log fold change, bubble color (green: males; magenta:hermaphrodites) represents enrichment in each sex. (C, E) Representative confocal micrographs showing multi-colored neuronal nuclei in the NeuroPAL strain otIs669 in head (C) and tail €where neurons are identified by colour barcode(*45*, *46*). These images were utilized to identify neurons expressing the *nlp-11(syb4759[nlp-11::SL2::GFP::H2B])* reporter in young adult hermaphrodites and males. Scale bars: 10µm. (D) Quantification of *nlp-11* fluorescent intensity from (C) in head neurons AUA, AVE, AIB URAD, URAV, M2, SABV, SIB AVM. (F) Quantification of *nlp-11* fluorescent intensity from (E) in tail neurons DVC, PDB, PLML, PLMR, and DVB neurons in both sexes at YA stage. n represents number of animals in each group in which neurons were identified and recorded. Number of worms (hermaphrodite, male) were: AUA, AVE, URAD, URAV, PDB, PLML, PLMR (14,14); AIB (12,12); M2 (12,13); SABV, AVM (13,13); SIB (9,9); DVB (14,12); DVC (14,13). Box-and-whiskers plots show median (center line), 25–75th percentiles (box), min– max (whiskers), with dots representing data points. Two-sided Mann-Whitney test was used for comparisons.

We validated the predicted sexual dimorphism in the expression pattern of two broadly expressed neuropeptide genes: *nlp-11* and *flp-20,* focusing on clusters within the head or tail regions. *nlp-11* showed dimorphic expression in 12 out of 16 predicted sex-shared neurons and in several male-specific neurons (Fig.3B-F, fig. S5A-B). The remaining four neuron types were either located in the midbody or were difficult to identify. Similarly, *flp-20* showed sexually dimorphic expression in four sex-shared neurons and in the male-specific CEMs (fig. S5C-G).

When examining our dataset, we wondered how many genes show a predicted binary dimorphic expression. To explore this notion, we analyzed binary gene expression (*See Methods*). All 62 sex-shared clusters showed many genes with dimorphic binary expression, supporting the existence of previously unrecognized sexually dimorphic neurons. Interestingly, some clusters showed a sex-bias gene expression profile, for example, SIA, where most of the binary genes were hermaphrodite-biased (Fig. 2E). Next, we asked how many of the DEGs we found in Fig. 2A are predicted to have binary expression using this approach and discovered that while some clusters have many DEGs that are also binary (such as AIM, BDU, and PHC), some have almost none (PVQ and RIV) (Fig. 2F). Furthermore, we calculated the number of predicted binary genes out of the total genes in our dataset and found that almost 70% of the genes in each sex across all cell clusters show binary expression, indicating how widespread sexually dimorphic gene expression is. Most of these genes are common to both sexes, indicating that different genes are differentially expressed according to the cell type, with around 1000 genes that are exclusively binary in each sex (across all cell clusters), supporting future on/off switches that could be discovered (Fig. 2G). Together, these findings reveal previously unrecognized sexually dimorphic neurons, highlight a broad landscape of sex-biased gene expression with a marked enrichment for neuropeptides, and indicate that predicted binary expression patterns are a prominent feature of this molecular dimorphism.

### A variety of neuronal gene families are differentially expressed between the sexes

In addition to the neuropeptide gene family, other gene families could point towards sex differences in neuronal function (fig. S4,6). A large fraction of synaptic-related genes and ion channels were differentially expressed (Fig. 2C, Fig. S4C-I). Many transcription factors (TFs) were cell-specifically differentially expressed between the sexes (fig. S6A). TFs known to drive sex-specific maturation, such as *tra-1* and *mab-3*(*15*), were differentially expressed in only a few neurons (fig. S6A-B) and not ubiquitously, perhaps because of the age of the animals used in our analysis. Previous data support a decrease in their expression after sexual maturation(*31*). The TF *tbx-32* was differentially expressed in the largest number of clusters with a male bias, hinting at a previously uncharacterized role for this gene in the development of the male nervous system (fig. S6A). DEGs also included TFs that are predicted to be bound by TRA-1 (*15*, *43*, *44*), supporting possible multiple unexplored roles for TFs in sex-specific neuronal regulation (fig. S6B).

### Sex differences in neurotransmitter gene expression shape neuronal output without altering identity

Neuronal terminal identity is defined by the expression of specific effector gene batteries, which are directly activated and maintained by transcription factors defined as ‘terminal selectors’(*47*, *48*). We compared the expression of known terminal selectors and found that most of them showed similar expression patterns, with few exceptions, for example *ast-1,* which is known to dictate CEP, ADE and AVG identities(*49–51*), is enriched in male SMD or *ceh-31,* which is known to dictate PVR identity(*47*), is enriched in male AVB (fig S6C). We next examined neurotransmitter-usage(*35*, *52–55*). For this purpose, we examined the expression of neurotransmitter-related genes in hermaphrodite and male sex-shared neuronal clusters (same clusters shown in Fig. 2A, Fig. 4A). While the known AIM neurotransmitter switch(*35*) is fully captured in our data set (Fig. 1F-G, Fig. 4A), we did not observe any other similar cases of neurotransmitter identity switch (Fig. 4A). However, when examining the differential expression of these neurotransmitter-related genes, we found that some of them, mostly cholinergic-related and glutamatergic-related genes, exhibited biased expression in several sex-shared clusters (Fig. 4B), suggesting sexual dimorphism may be more prominent in neurotransmitter release and synthesis rather than identity.

**Fig. 4.**
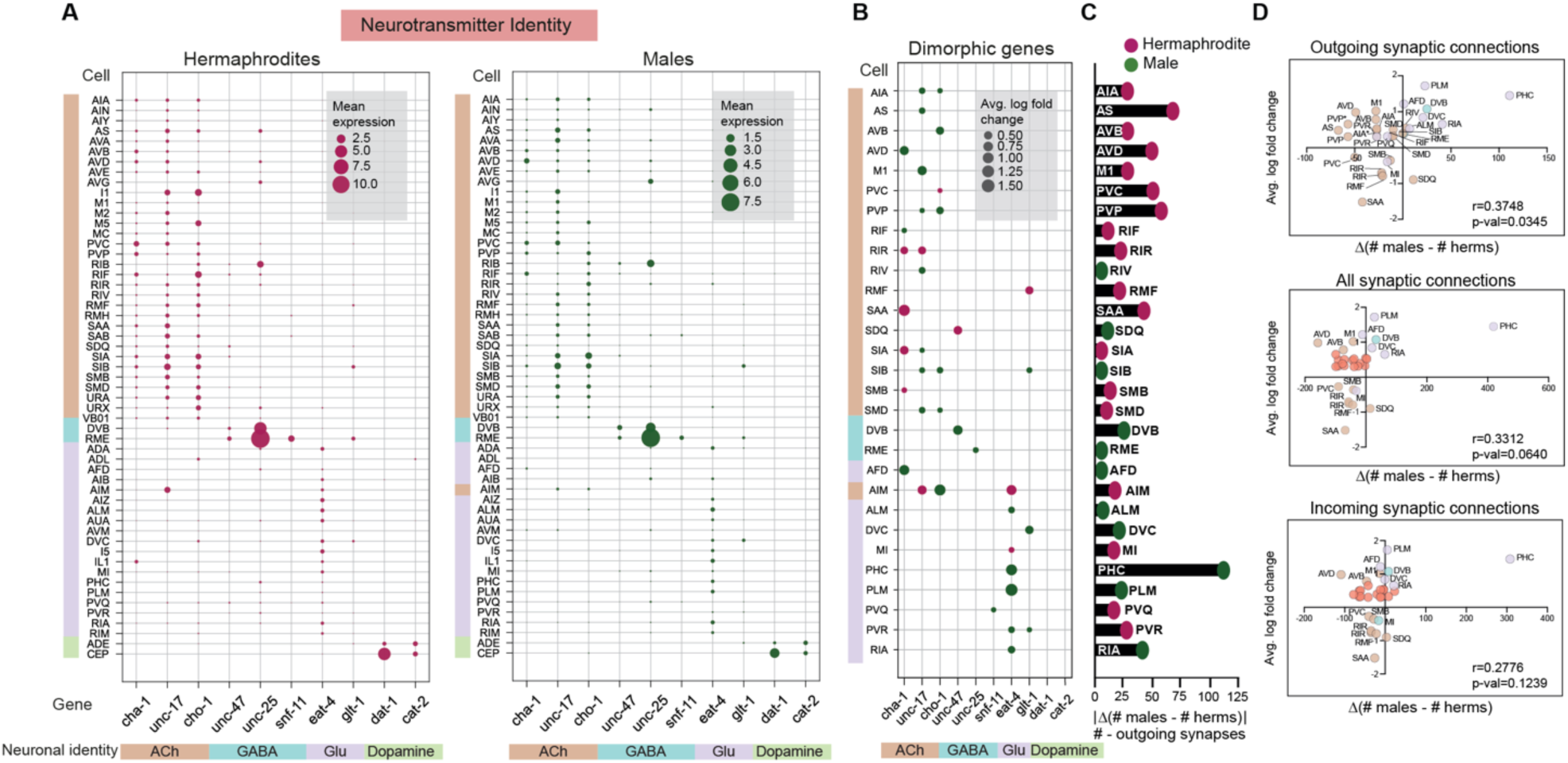
Neurotransmitter identity is preserved across sex-shared neuronal clusters. (A) Left and right bubble plots representing the mean expression values of neuronal identity genes in hermaphrodites and males, respectively. Neuronal clusters and genes are color-coded according to their neurotransmitter identity. (B) Bubble plot representing differentially expressed neurotransmitter identity genes between the sexes. Only genes with significant log fold change when comparing the hermaphrodite and male mean expressions are shown. Bubble size in (A) corresponds to gene expression and in (B) to the average log fold change in gene expression. Bubble color indicates enrichment in either sex (green, male; magenta, hermaphrodite). Rows represent neuron types, grouped color coding for neurotransmitter identity. Columns represent individual genes. (C) Bar plot representing the difference in outgoing connectivity between males and hermaphrodites in each cluster (The number of outgoing connections in males minus the number of outgoing connections in hermaphrodites for each neuron that was examined). Numbers are represented as an absolute value (*See Methods* for more details). Magenta represents a higher value in hermaphrodites, while green represents a higher value in males. (D) Two-tailed Pearson correlation analysis between the difference in connectivity (outgoing connections, all synaptic connections, and incoming connections) and the average log-fold change of neurotransmitter identity genes (*See Methods* for more details). The numbers representing log fold change were correlated with the number representing the difference in outgoing connectivity. The colors of the dots correspond to the neurotransmitter identity colors shown, except those that overlap and are labeled red (A). For (D) we performed a two-sided Pearson correlation test.

With this in mind, we explored the relationship between dimorphic neurotransmitter-related gene expression and dimorphic synaptic connectivity. The complete synaptic wiring diagram of the nervous system is available in *C. elegans* for both sexes(*37*), enabling a comparison of the number of synaptic connections to neurotransmitter-related DEGs. We hypothesized that neurons that show higher expression of neurotransmitter-related genes in one sex, might also have more outgoing connections. To test this, we correlated the difference in connectivity between the sexes in outgoing, incoming, and all synaptic connections (chemical connections)(*37*) (Fig. 4C*, See Methods*) with the average log-fold change of neurotransmitter-related genes presented in Fig. 4B. Strikingly, Pearson correlation coefficient was significant only for outgoing connections but not for incoming or all synaptic connections (Fig. 4D). For example, sex specific enrichment of the acetyltransferase *cha-1*, which is required for acetylcholine synthesis(*56*), was observed in the sex-shared neurons SIA, SAA, RIR, and AFD, aligning with increased connectivity of these neurons in the respective sex. Conversely, sex specific enrichment of the vesicular glutamate transporter *eat-4*(*57*) corresponded with greater connectivity in the neurons RIA, PLM, PHC, ALM, and AIM in the respective sex. (Fig. 4B-C). Thus, to some extent, neurotransmitter release of a neuron is correlated with its outgoing connectivity.

Altogether, our findings suggest that while neurotransmitter identities are largely preserved in sex-shared neurons, sexually dimorphic expression of neurotransmitter-related genes correlates with differences in outgoing synaptic connectivity, pointing to a mechanism by which functional dimorphism may arise without altering core neuronal identities.

### Sexually dimorphic gene expression and function of the PLM neuron underlie sex-specific mechanosensory responses

Among the sex-shared neuronal clusters with a high number of DEGs, we observed all the touch receptor neurons (TRNs): ALM, AVM, PVM, and PLM (Fig. 2A), required for mechanosensory responses to gentle touch(*58*). We turned to explore this dimorphic molecular repertoire, focusing specifically on PLM, a TRN that mediates the posterior gentle touch response(*58*) and is the most sexually dimorphic among TRNs (Fig. 2A, 5A)(*58*, *59*). We validated the differential expression of multiple neuronal-related genes (Fig. 5B-C) using transcriptional and translational reporters (Fig. 5D-G), or single molecule Fluorescent In Situ Hybridization (smFISH) (Fig. 5H). We tested the gentle touch responses of hermaphrodites and males. We found that hermaphrodites showed reduced responses to posterior but not to anterior gentle touch (Fig. 5I). As a control, we tested harsh touch-tail responses in the same animals, which were not dimorphic (Fig. 5I). While characterizing the development and growth of PLM, we found sex differences in the length of the posterior processes at the adult stage, with fewer males having posterior processes (fig. S7A-D). Analysis of PLM connectivity showed minor differences between the sexes, with males making slightly more gap junctions(*37*) (fig. S7E). Thus, we speculated that the behavioral dimorphism we observed in posterior gentle touch could stem from sex differences in PLM neuronal activity. Indeed, when measuring calcium traces of PLM basal activity, we observed higher activity in males in frequencies up to 2 Hz (Fig. 5J-M), further supporting a sexually dimorphic role for PLM. In addition, optogenetic activation of the tail TRNs, which includes PLM, resulted in male behavioral responses that were statistically significantly distinct from hermaphrodites, supporting PLM dimorphism (fig. S8). We did not observe significant changes in the expression of major touch-response genes such as *mec-4* and *mec-10*(*60*, *61*) (table S3). To test the involvement of specific DEGs in PLM function, we knocked down the neuropeptide *nlp-11* (higher in PLMs of males) or the RNA-binding protein *mec-8* (higher in hermaphrodites) and assayed the RNAi-silenced animals or mutant animals for posterior gentle touch responses in both sexes. While *nlp-11* did not show an effect on gentle touch responses in either of the sexes, *mec-8* was required only in hermaphrodites (Fig. 5N). Importantly, over-expressing MEC-8, specifically in PLM, restored the behavioral responses of hermaphrodites (Fig. 5O). Altogether, our results establish PLM as a sexually dimorphic neuron with a differential transcriptomic profile and identify a dimorphic gene expression critical for a sex-specific mechanosensory response.

**Fig. 5:**
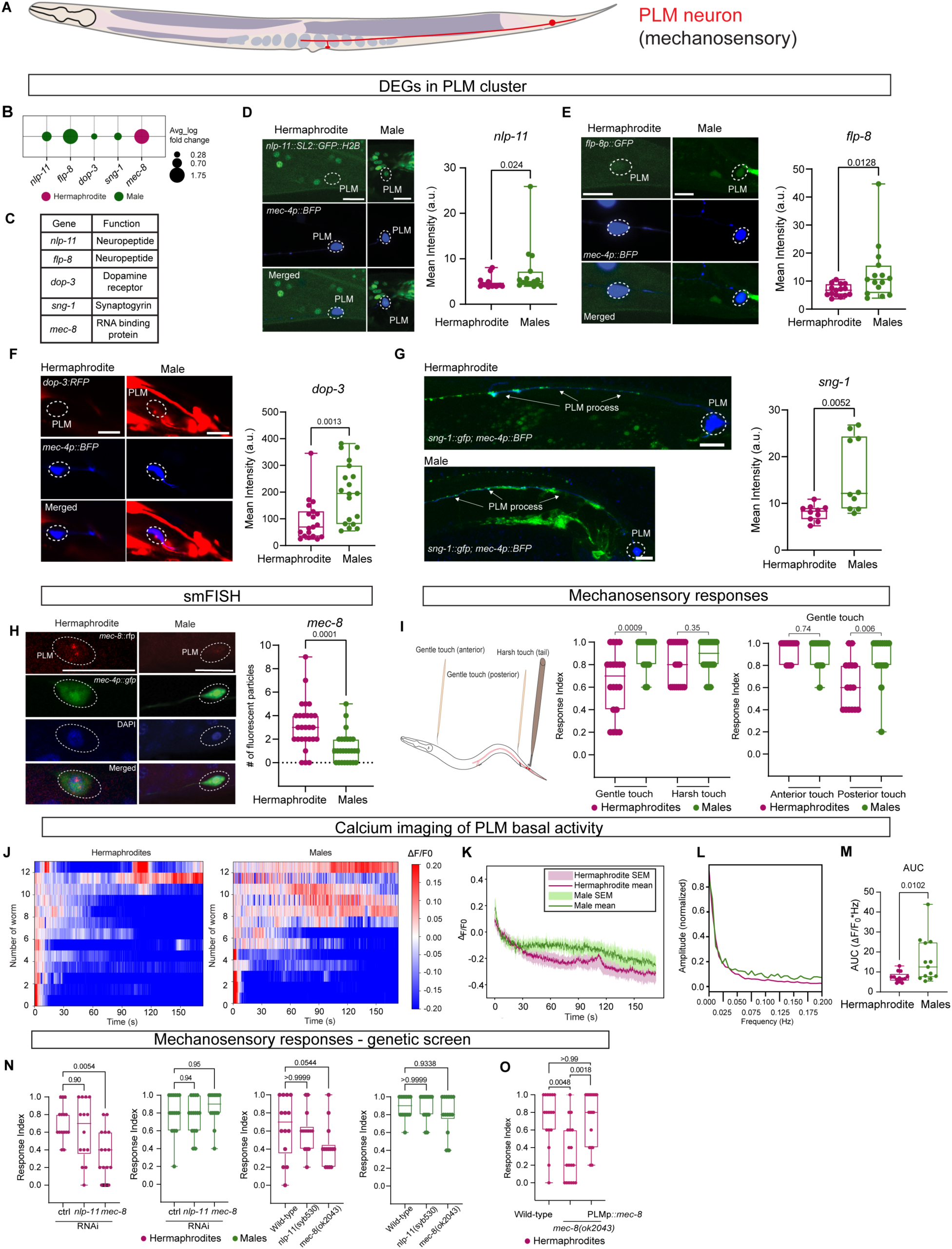
Sexually dimorphic properties of PLM neuron. (A) Illustration of PLM sensory neuron (https://www.wormatlas.org/) (B) Bubble plot of *nlp-11, flp-8, dop-3, sng-1,* and *mec-8* expression in PLM. Size: average log fold change; color: green: male and magenta: hermaphrodite enrichment. (C) Table showing gene functions from (B). (D-H) Representative confocal micrographs and quantification of gene expression in PLM (white dotted circle): (D) *nlp-11::SL2::GFP::H2B* (n=16 hermaphrodites, 15 males) (E) *flp-8p::GFP* (n=15/group) (F) *dop-3::RFP* (n=19/group) (G) *sng-1::GFP* in PLM neurite (n=10/group) (H) mec-8 smFISH probe with *mec-4::gfp* and DAPI (n=27 hermaphrodites, 26 males). Scale bars = 10 µm. a.u. = arbitrary units. (I) Left: Illustration of gentle/harsh touch in *C. elegans.* Created in BioRender. Oren, M. (2025) https://BioRender.com/i2zd1il Middle: posterior gentle touch responses of both sexes and harsh tail-touch responses tested on the same animals (n = 20/group). Right: anterior and posterior gentle touch responses of both sexes (n=15/group). (J) Normalized GCaMP6s calcium responses in PLM (n=13 animals per group). Heatmaps show calcium levels of individual worms. (K) Mean and SEM calcium traces from (J). (L) Mean traces showing normalized amplitudes of PLM, zoomed in at frequencies 0-0.2 Hz. (M) AUC (area under the curve) measured from PLM amplitudes for chosen frequencies (0-2 Hz). (N) Posterior gentle touch responses of both sexes after RNAi: control (n=15 each), *nlp-11* (n=14 hermaphrodites, 15 males), *mec-8* (n=15 hermaphrodites, 14 males), and mutants *nlp-11(syb530)*, *mec-8(ok2043)* (n=14 each). (O) Posterior gentle touch responses of hermaphrodites in Wild-type (n= 17), *mec-8 (ok2043)*(n=18), and *mec-8* rescue in PLM (n=18). Boxplots: median (line), 25th–75th percentile (box), min–max (whiskers), dots = individual data points. In (D, E, F, G, H, I, M) we performed a two-sided Mann-Whitney test for each comparison. In (N, O) we performed a Kruskal-Wallis test followed by a Dunn’s multiple comparison test.

### The cell adhesion molecule LAD-2 is required for proper neurite extension in male but not hermaphrodite PHC neurons

Our study offers molecular candidates to explain the mechanistic basis for previously recognized sexual dimorphism of neurons such as PHC, AFD, AIM and AVG (Fig. 2A). For example, we validated two differentially expressed neuropeptides, *flp-7* and *flp-22*, with higher expression in hermaphrodite AVG neurons, suggesting a potential role in modulating AVG’s circuit, specifically in hermaphrodites (fig. S9A-D)(*7*). We next focused on PHC tail sensory neurons, which undergo massive remodeling in males compared to hermaphrodites, with males exhibiting neurite extensions that are twice as long(*36*). This elongation is presumably required to accommodate the increased synaptic connectivity with male-specific neurons(*36*, *37*) (Fig. 6A, Fig. 4D). The TF DMD-3 was shown to modify the function, connectivity, and molecular features of the male PHC neurons. We observed increased expression of *dmd-3*, *eat-4/VGLUT* and the neuropeptide *flp-11* in the male PHC cluster(*34, 36*) (Fig. 6B). Our ranking analysis identified 209 DEGs for PHC (13 in male neurons and 16 in hermaphrodite neurons, table S2, Fig. 2A). Supporting the dimorphic connectivity and morphology of PHC, these DEGs were enriched in the GO-terms “synaptic function” and “transmembrane transport” (Fig. 6C). We validated a subset of neuronal DEGs predicted by our scRNA-seq using transcriptional and translational reporters, including a neuropeptide gene, GABA and dopamine receptors, and the cell adhesion molecule *lad-2* (Fig. 6D-I). Since *lad-*2 has been shown to mediate axon guidance(*62*), we imaged the neurite extension of PHC in both sexes in *lad-2* mutants. We revealed that *lad-2* is required only for the male extension, consistent with its higher expression in males (Fig. 6I-K). Taken together, our data set confirms the dimorphic transcriptomic profile for known dimorphic sex-shared neurons and allows the identification and exploration of previously unrecognized molecular mechanisms within these neurons.

**Fig. 6:**
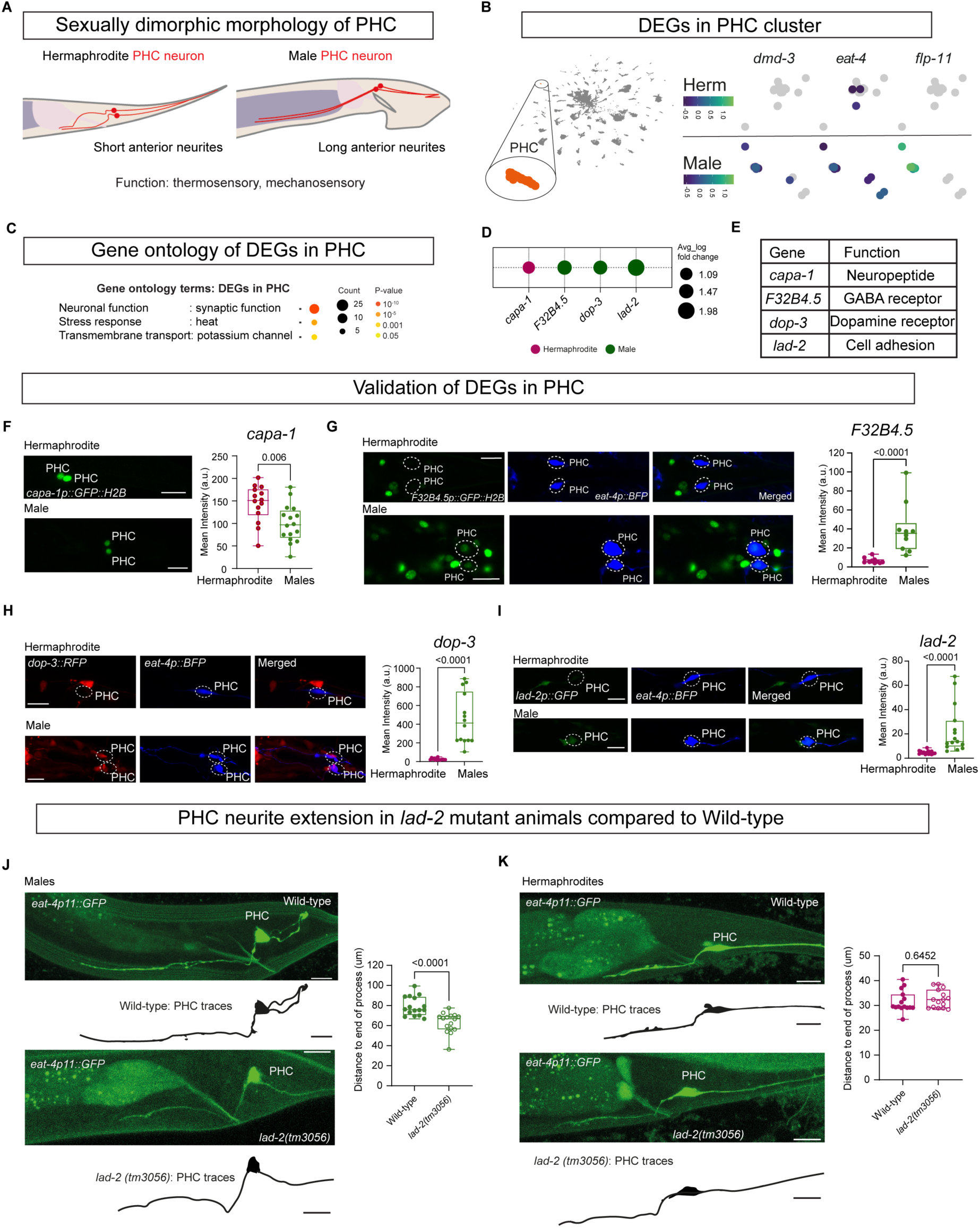
Expression of DEGs in PHC. (A) Illustration depicting the location of PHC sensory neurons and their neurite length in both sexes, as outlined in WormAtlas. Created in BioRender. Oren, M. (2025) https://BioRender.com/i2zd1il. (B) Validation of previously reported DEGs. Left, PHC representation in the UMAP projection of all cells (inset marked as orange). UMAP projection depicting expression of *dmd-3*, *eat-4*, and *flp-11* gene within PHC cluster. Heatmap represents expression level. (C) GO term enrichments for DEGs in PHC cluster, including the number of genes in each GO term. (D) Bubble plot representation of *capa-1, F32B4.5, dop-3,* and *lad-2* differential genes expression in PHC cluster. Bubble size represents average log fold change, and bubble color (green: males and magenta: hermaphrodites) represents enrichment in each sex. (E) Table lists the established functions for genes in (D). (F-I) Representative confocal micrographs showing expression and quantification of various genes in PHC neurons (white dotted circle). (F) *capa-1p::GFP*. Hermaphrodites, n = 14; males, n= 16. (G) *F32B4.5p::GFP,*n = 10 animals per group. (H) *dop-3p::RFP*. n = 14 animals per group. (I) *lad-2p::GFP,* n = 15 animals per group. (J-K) Representative confocal micrographs of *eat-4p11::GFP* expression in PHC neurons in wild-type and *lad-2 (tm3056)* mutant males, n = 16 animals per group (J) and hermaphrodites, n = 10 animals per group (K). Scale bars represent 10 µm. In the box-and-whiskers graph, the center line in the box denotes the median, while the box contains the 25th to 75th percentiles of the dataset, whiskers define the minimum and maximum value with dots showing all points. In (F, G, H, I, J, K) We performed a two-sided Mann-Whitney test for each comparison.

### Correlation analysis predicts regulators of synaptic connectivity

Can we predict which genes regulate synaptic connectivity by analyzing their expression patterns in relation to wiring maps? Understanding the link between gene expression and synaptic connectivity is a key question in neurobiology. By integrating transcriptomic profiles with connectome maps, we can generate predictions on the identity of genes that regulate synaptic wiring. While previous studies have been limited to hermaphrodites and relied on binary gene expression data, leaving room for broader and more nuanced investigations(*63*, *64*). As the connectome of *C. elegans* is available for both sexes, we further utilized our non-binary single-cell gene expression dataset of both sexes and the connectome of *C. elegans* to answer this key question. For this purpose, we calculated the correlation between the number of outgoing connections for each neuron based on(*37*) and the average gene expression values of all genes in our dataset, across the 62 clusters ranked in Fig. 2A, analyzing hermaphrodites and males separately (*See Methods*, Fig. 7A-B). We chose outgoing connectivity because of the availability of pre-synaptic *in-vivo* markers(*65*, *66*) for further validation of candidates. To identify genes that might regulate the outward connectivity of neurons, we performed statistical analysis accounting for multiple comparisons, which resulted in an adjusted p-value for each correlation value (*See Methods*). We then searched for significantly correlated genes that overlap between hermaphrodite and male correlation profiles (Fig. 7C). We discovered 54 genes with significant correlation in both sexes, with more than half still uncharacterized. Among the neuronal genes exhibiting a significant positive correlation, are key presynaptic genes such as the active zone scaffold *syd-2* involved in presynaptic assembly(*67*, *68*), but also postsynaptic genes such as *acc-2,* encoding acetylcholine-gated chloride channel(*69*), and *gar-3* encoding G-protein-linked acetylcholine receptor(*70*) (Fig. 7C). This finding highlights numerous uncharacterized genes with potential synaptic function and suggests that some synaptic regulatory pathways are shared between the two sexes. To gain a better perspective into dimorphic synaptic regulation, we looked specifically for neuronal genes that are either significantly correlated with outgoing connectivity in hermaphrodites, in males or in both sexes (Fig. 7D). While exploring the expression of these genes in our data set, we uncovered the sexually dimorphic expression patterns of some of them (Fig. 7E). Importantly, these genes included *ttx-7,* which encodes a myo-inositol monophosphatase (IMPase), already shown to mediate proper synaptic localization in the interneuron RIA(*71*) and *zig-1* predicted to encode a cell adhesion molecule whose function is unexplored(*72*, *73*). We found that *ttx-7* is significantly correlated in both sexes (Fig. 7C). Based on *ttx-7* expression profile, we hypothesized that it might regulate synaptic connectivity of RIA neurons in both sexes (since both express it equally high), but its absence would have a greater effect on the synaptic connectivity of hermaphrodites CEP neurons (since it is highly expressed in hermaphrodites) (Fig. 7E, red squares, and 7F). We indeed found that in *ttx-7* mutant animals, RIA pre-synapses were mostly mislocalized in both sexes (Fig. 7G), which is partly explained by elongated RIA processes observed in some of the animals (fig. S9E-F), while the number of pre-synapses in CEP neurons was reduced only in hermaphrodites (Fig. 7I). *zig-1* mutant animals showed lower pre-synaptic signal only in males, in concordance with the gene being significantly correlated only in males (Fig. 7D, Fig. 7H). In summary, combining our dataset together with connectivity maps uncovers genes regulating synaptic wiring, both in non-dimorphic and dimorphic contexts.

**Fig. 7.**
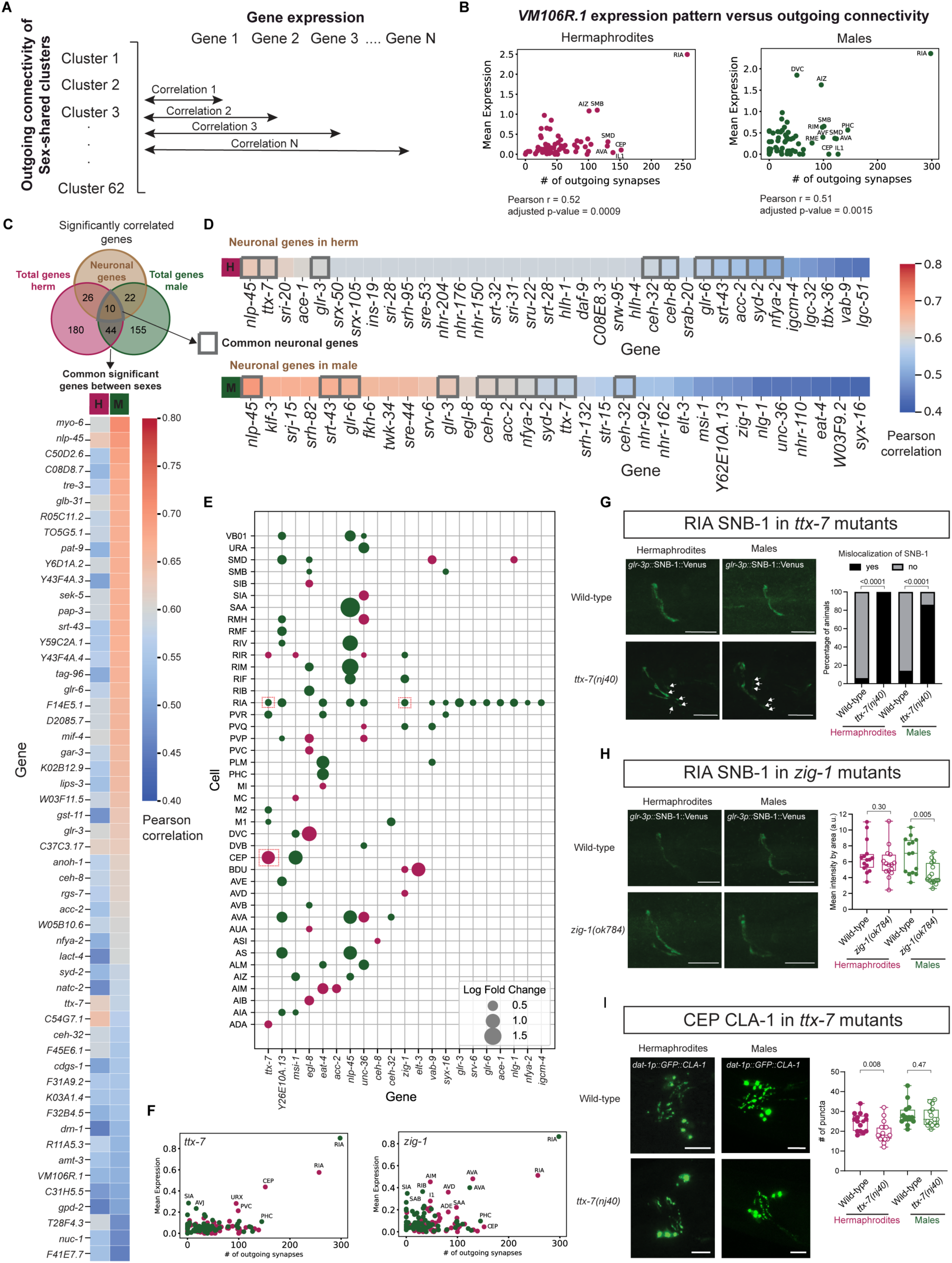
Identification of genes regulating connectivity. (A) Schematic of correlation analysis. Rows: number of outgoing connections calculated according to(*37*) across neurons. Columns: mean expression of genes across neurons. Pearson correlation was calculated for connectivity numbers and mean expression per gene, separately for hermaphrodites and males. (B) Mean expression of *VM106R.1* across outgoing connections in both sexes. (C) Venn diagram showing significantly correlated genes in hermaphrodites and in males and the number of neuronal genes within these groups (brown) (top). Heatmap of Pearson correlation coefficients of significantly correlated and overlapping genes (bottom). (D) Heatmaps of Pearson correlation coefficient of significantly correlated neuronal genes in hermaphrodites (top) and in males (bottom). (E) Expression of significant neuronal genes from (D). Size represents average log fold change, color represents sex enrichment. (F) Mean expression of *zig-1* (right) and *ttx-7* (left) across outgoing connections in hermaphrodites and males. (G-I) Representative confocal micrographs and quantification of: (G) *glr-3p::SNB-1::venus* in RIA in Wild-type (hermaphrodites n = 15, males n = 14) and *ttx-7(nj40)* (n = 15/group) (left). Percentage of animals showing mislocalization of SNB-1 (right). (H) *glr-3p::SNB-1::venus* expression in RIA in Wild-type (hermaphrodites n = 15, males n = 14) and *zig-1(ok784)*(n = 15/group)(left). Quantification of venus signal (right). (I) *dat-1p::novogfp::CLA-1* expression in CEP neurons in Wild-type and *ttx-7(nj40)*(left). Quantification of CLA-1 puncta in CEP (right). n = 15/group. In the box-and-whiskers graph, the center line in the box denotes the median, while the box contains the 25th to 75th percentiles of the dataset, whiskers define the minimum and maximum value with dots showing all points. (C) Pearson correlation analysis with Benjamini Hochberg correction for multiple comparisons was done. (G) Two-sided Fisher’s exact test was done for each comparison. (H-I) Two-sided Mann-Whitney test was done for each comparison. Magenta: hermaphrodites, green: males.

## Discussion

In this work, we established a transcriptome map of the sex-shared nervous system in both sexes and uncovered previously unexplored sexually dimorphic neurons, including the posterior light touch neurons PLM. In addition to sex-shared neurons, *C. elegans* males have an additional 93 male-specific neurons. Even though these can be subdivided into neuronal subtypes(*54*, *74*), for most male-specific neurons, molecular markers that define individual identities haven’t been described. Thus, male-specific clusters have been left out of this analysis until more tools become available(*75*). Our ranking analysis reveals that the most prominent dimorphic gene family is neuropeptides, with a male-biased expression, in accordance with our previously published developmental comparison between the sexes(*31*). Neuropeptides are short protein ligands that modulate synaptic activity mostly through the binding to G protein-coupled receptors (GPCRs), playing a critical role in regulating functions of the nervous system functions(*76*). A neuropeptidergic connectome has recently been constructed, revealing a dense network with unique sex-shared neurons that serve as “peptidergic hubs”(*77*). This suggests that a prominent rewiring of the neuropeptidergic connectome occurs in addition to the rewiring of the physical connectome between sexes. Future studies exploring the entire neuropeptidergic connectome in sexually dimorphic context would shed more light on how neuropeptide signaling has diverged to drive nervous system function and behavior. (*31*, *78*). Binary analysis of gene expression reinforced the strong dimorphisms identified in the ranking analysis and predicted approximately 1,000 sex-specific genes in each sex. These dimorphisms do not alter neuronal identity as terminal selector usage and neurotransmitter identity are mostly unchanged, suggesting that functional dimorphism may arise without altering core neuronal identities. Dimorphic neuromodulators and signaling molecules seem to be the major players that fine-tune sex-specific synaptic outputs. The few exceptions we found for terminal selectors predicted to be expressed in previously unidentified cell types might suggest that they play an active, sexually dimorphic role in shaping neuronal identity, or that their role is activity-related rather than identity-related.

Our functional analysis of sex-shared neurons uncovers previously unrecognized sexual dimorphism in mechanosensory posterior gentle touch responses, with males displaying higher sensitivity compared to hermaphrodites. This difference is likely mediated by the highly dimorphic touch receptor neuron PLM, which exhibits substantial differences in basal neuronal activity between the sexes. While PLM activity decreases when courtship begins(*79*), the heightened mechanosensory responses in males may be essential for mating, as was already shown for turning behavior (*80*) potentially through the neuropeptides *flp-8* and *flp-20*, which are enriched in male PLMs^73^. Hermaphrodites, on the other hand, do not display such behavior and show lower sensitivity. In contrast, both sexes exhibit similar responses to harsh tail touch(*7*), suggesting that hermaphrodites maintain robust responses to dangerous stimuli for survival. Our focus on mechanosensation in this and previous work(*7*) provides insights into how different touch modalities (gentle and harsh) can result in distinct outcomes, potentially reflecting different evolutionary pressures.

We identified the RNA binding protein MEC-8 as a hermaphrodite-enriched gene in PLM and showed it to be critical for their PLM mechanosensory responses, suggesting that different molecular mechanisms mediate mechanosensation in the two sexes. For example, *inx-18* and *eat-4* are enriched in male PLMs, pointing towards enhanced regulation of their post-synaptic partners, electrical and chemical, respectively. These genes may regulate the increased number of connections in male PLMs(*37*), possibly contributing to the robust touch response.

Our single-cell data set identified the cell adhesion molecule LAD-2 as a male-specific regulator of PHC neurite extension(*36*), though which specific isoform is responsible remains unclear(*62*). Future screens are required to uncover genes governing PHC neurite structure in hermaphrodites. We observed that neurotransmitter identity is largely preserved across sexes, with the notable exception of AIM neurons(*35*). Slight changes in expression levels for neurotransmitter genes between the sexes correlate with changes in outgoing connectivity, although this correlation was relatively low (r=0.37), suggesting other factors might underlie the differences in outgoing connectivity, such as synaptic-related genes, transcription factors, or a combination between different genes. This led us to explore the correlation between gene expression across our data set and connectivity. Based on this, we were able to predict a large number of candidate genes in both sexes that might regulate outgoing connectivity, validating two of them. While this correlation analysis was sufficient to identify gene-to-connectivity matches for specific neurons, it could not be extended to the entire worm’s nervous system, as our analysis did not capture all sex-shared neurons (62 clusters that have passed all the filters in ranking analysis) and some of the sex-shared neurons are under-represented in our data set. Further studies profiling the rest of the sex-shared neurons will possibly allow for a more accurate prediction of the way gene expression regulates connectivity. Even with a full transcriptomic representation of the nervous system, predictions can be complicated due to interactions existing between genes and to differences between connectivity and single-cell RNA sequencing data pooling (small sample size in EM sections versus pooling large number of worms for single-cell).

Together, our findings reveal that sexual dimorphism in the nervous system can arise through widespread modulation of gene expression, particularly among neuromodulators and signaling molecules, without altering core neuronal identities. By linking these molecular differences to synaptic connectivity and behavior, our work uncovers a flexible regulatory layer that tunes shared circuits for sex-specific function by linking these molecular differences to synaptic connectivity.

## Materials and Methods

### *C. elegans* strains and maintenance

All *C. elegans* strains used in this study were cultivated according to standard methods(*81*) and listed in table S10. Wild-type strains were Bristol, N2. *him-5(e1490)* or *him-8(e1489)* were treated as controls for strains with these alleles in their background. Worms were grown on nematode growth media (NGM) plates seeded with *E. coli* OP50/NA22 bacteria as a food source. Worms were raised at 20°C, unless noted otherwise. The sex and age of the animals used in each experiment are noted in the associated figures and legends. All transgenic strains used in this study are listed in the key resources table.

### Isolation of *C. elegans* pure male population

To generate pure male populations, we followed our previously described protocol(*31*). Briefly, *C. elegans* strain with the genotype *rab-3p(prom1)::2xNLS::TagRFP] IV; dpy-28(y1) III; him-8(e1489) IV* was employed. This strain harbors a temperature-sensitive mutation (*y1*) in the dosage compensation gene, *dpy-28* along with *him-8(e1489)* mutation and a pan-neuronal RFP. Worms were cultured at 15°C on 15 cm NGM plates seeded with NA22 bacteria. Embryos were isolated by hypochlorite treatment and were age-synchronised 14-16 hours on a foodless NGM plate at 25°C. L1 larvae were collected and were placed on a food NGM plates approximately 1.5 cm from the food on NGM plates and were allowed to crawl towards food for 1-2 hours at RT. Using a scalpel, the area where L1/embryos were initially placed was removed. L1 worms were collected and washed with M9 buffer and were incubated at 20°C until adulthood. This resulted in male population of ≥98% purity. For each experiment the percentage of males in the population was recorded.

### Worm single cell dissociation

Worms of the desired genotype were grown at 15°C on 15 cm NGM plates seeded with NA22 bacteria until the plates were populated. Gravid adult worms were collected in M9 buffer and embryos were isolated by hypochlorite treatment. The obtained embryos were hatched for 14-16 hours on a foodless NGM plate at a restrictive temperature of 25°C. The number of L1 larvae were counted and were plated (40,000-50,000 animals per plate) on a fresh NA22 seeded bacterial plates and were grown to young adult stage at 20°C. Young adult worms were washed 10 times at 1300 rpm for 2 minutes with M9 buffer to get rid of any adhering bacteria. The worms were then washed with ddH_2_O. The worm pellets were then subjected to single cell dissociation using protocols described earlier with slight modifications(*20*, *82–85*). For every 0.5 ml of worm pellet 1 ml of freshly thawed sterile SDS-DTT solution (200 mM DTT, 0.25% SDS, 20 mM HEPES, 3% sucrose, pH 8.0, thawed freshly) was added and incubated at room temperature for 4 min with gentle mixing every 1 minute. Immediately after SDS-DTT treatment, worms were pelleted at 3,000 rpm for 1 min and washed 5 times with 10 ml egg buffer (118 mM NaCl, 48 mM KCl, 2 mM CaCl_2_, 2 mM MgCl_2_, 25 mM HEPES, pH 7.3). 0.5 ml aliquots of worm pellet were distributed to 1.6 mL microcentrifuge tubes. 1 mL freshly prepared 15 mg/mL Pronase-E was added to each tube and flicked to mix and then rotated at 20°C for 5 minutes. Using a glass pasteur pipette, worm suspensions were combined from multiple tubes and were transferred to a 7 mL dounce homogenizer (pestle clearance 0.02−0.056 mm, Kimble Chase). Pronase E was added to a final ratio of 4:1 relative to the worm pellet. If the worm pellet was smaller than 0.5 mL a 2mL glass homogenizer was used. The animals were disintegrated to single cells using type “B” plunger for 4-5 minutes at room temperature and then continued on ice gently. We note that homogenizing on ice increased cell survival. The status of dissociation was monitored under a dissecting microscope at 5-minute intervals. Dissociation was stopped when most of the animals were dissociated but a few whole or fragmented animals were still present. Using a glass pasteur pipette, the cell suspension was aliquoted into 1.6 mL microcentrifuge tubes that were centrifuged at 5,000 x g for 5 minutes at 4 °C. Supernatant was removed and the cell pellet was washed 3 times with 1 mL ice-cold egg buffer per wash at 5,000 x g for 5 minutes at 4 °C. The cells pellet was resuspended in 1 ml ice-cold egg buffer and was passed through 5 μm syringe filter gently. Cell suspension was taken on ice to FACS sorting immediately.

### FACS based sorting

Single cell suspension from the desired strain were subjected to Fluorescence Activated Cell Sorting (FACS) on a BD FACSAria™ III Cell Sorter using a 70-micron nozzle. eFlour 450 dye (final concentration of 1 μl/mL) was used to discriminate live and dead cells. The cells were gated for side scatter (SSC)-Area vs forward scatter (FSC)-Area, then single cells were selected from doublet using FSC-Height vs FSC-Area. Finally, RFP+ cells were gated and sorted under the “4-way Purity” mask from this subpopulation. Wild-type *him-5(e1490)* without any RPF+ cells was used as control for gating. Sorted cells were collected in 1.6 mL microcentrifuge tube containing 500 µl L-15-20 (L-15 medium containing 20% fetal bovine serum). The suspension was centrifuged at 5,000 x g for 10 minutes at 4 °C, resuspended in 50-80 µl of L-15-20 and counted using a hemocytometer.

### RT-PCR

For carrying RT-PCR experiments a previously described protocol was used(*31*). YA hermaphrodite worms were dissociated into single cells and the RFP+ and RFP-cells were FACS sorted in 500 µL Trizol, flash frozen in liquid nitrogen and stored at −80°C until further processing. Total RNA was extracted using the TRIzol LS (Invitrogen) protocol until the isopropanol precipitation step then was re-suspended in the extraction buffer from RNA isolation kit (PicoPure, Arcturus). The manufacturer’s recommendations were followed for further isolation. Using the SuperScript^TM^ IV first-strand synthesis system (Invitrogen, Catalog # 18091050) and random hexamers, the extracted total RNA was then converted to cDNA. Fast SYBR™ Green Master Mix (Applied Biosystems, Catalog # 4385612) was used to carry out qPCR as per manufacturer protocol. The experiments were carried out in 3 biological replicates. *act-1, pmp-3* and *tba-1* genes served as internal controls. Ct values were obtained, and relative quantification of the expression of the target genes was performed utilizing 2−ΔΔCt method.

### Cell imaging (imaging flow cytometer)

Cell imaging experiments were carried out using an ImageStreamX Flow Imager system to evaluate the morphology and viability of cells after single cell dissociation. Single cells were dissociated from a *C. elegans* strain expressing pan-neuronal TagRFP as described above and were resuspended in PBS with eFlour 450 dye (Live/Dead) and DRAQ5 dye (DNA binding) to a final concentration of 1 μl/mL. The positive samples containing all fluorochromes were used to set up each channel and the laser intensities. Gating was conducted based on Area vs RFP intensity, followed by the selection of RFP+ cells and by discriminating fluorophore using Area vs intensity. Cells derived from Wild-type N2 without RFP+ expression served as a control for gating. Images were acquired for 20,000 cells.

### Single-cell RNA sequencing

Depending upon the cell concentration, the single-cell suspensions were diluted to reach the optimal range as recommended by 10x genomics manufacturer’s protocol (approx. 800-1000 cells/μl). Each lane in the 10x genomics chip was targeted for 8,000 or 10,000 cells. Libraries were prepared with 10x Genomics v3 Chemistry according to the manufacturer’s protocol. The libraries were sequenced using the Illumina NovaSeq 6000 with 150 bp paired end reads, except for one hermaphrodite sample which was sequenced using the Illumina Nextseq 500 with a 75 cycle high output kit and paired end sequencing. Detailed experimental information is found in table S4. Real-Time Analysis software (RTA, version 4.0.0; Illumina) was used for base calling and analysis was completed using 10X Genomics Cell Ranger software (4.0.0). Detailed experimental information is found in table S4.

### Single-cell RNA-Seq Mapping

Sequenced reads from each experiment were mapped to the *C. elegans* reference transcriptome, version [WBcel235](https://www.ncbi.nlm.nih.gov/datasets/genome/GCF_000002985.6/) release WS276, using the Cell Ranger pipeline version 4.0.0. We extended the 3′ UTR of each gene by 500 base pairs for additional mapping of reads to the 3′ extremity of the gene body since the 3’ end is poorly annotated in *C. elegans*(*26*).

### Background correction and filtering procedure

For each experimental batch (6 hermaphrodite and 6 male samples), the empty droplets were identified, and background correction was performed using SoupX(*86*) as previously described(*20*). Each batch of cells was filtered by selecting clusters based on neuronal and non-neuronal gene content as detailed below (fig. S10A). First, UMAP dimensional reduction(*87*) was performed using 50 PCs in Seurat(*33*) V3.2.2. Later, clusters were defined by computing Shared Nearest Neighbor using two UMAP dimensions and the original Louvain algorithm with a resolution of 1, using FindNeighbors and FindCluster functions where all other parameters were set to default. Clusters with more than 0.5% neuronal markers and less than 1.5% non-neuronal markers were selected for further analysis (fig. S10B) using a list of 563 neuronal and 114 non-neuronal marker genes based on table S1. To ensure no loss of neurons through this procedure, a second round of selection was performed on cells that did not pass the first filtering step. For the second round, subclustering was performed on the remaining clusters using same parameters as before and the same filtering of clusters using neuronal and non-neuronal markers was applied. Finally, cells containing more than 20% of UMIs originating from mitochondrial genes were removed (fig. S10D-E).

### Dimensionality reduction, batch correction and cluster Identification

Monocle 3(*88*) was used to combine the male and female batches with the CeNGEN(*20*) L4 hermaphrodite neuronal data. Counts were imported to Monocle 3 and batches were aligned using Batchelor(*89*) using 135 PCs and an alignment k of 5, followed by UMAP dimensional reduction using a minimal distance of 0.3 and 75 neighbors, and clustering using Leiden algorithm with a resolution of 3e-3, resulting in 72 partitions. This UMAP projection was kept to display the final clustering and annotation analysis. Two iterative rounds of subclustering were performed after the first one. For the second iteration 2 subclustering steps were performed on partition 1 and on the merge of partitions 2 to 72. The same parameters were used for both subclusterings: 135 PCs and an alignment k of 20, followed by UMAP dimensional reduction using a minimal distance of 0.1 and 15 neighbors, and clustering using Leiden algorithm with a resolution of 3e-4 resulting in 36 and 82 partitions, coming from former partitions 1 and 2-72, respectively. This totaled 118 partitions that were inspected to assign identities or to continue with a third round of subclustering. We used the content of CeNGEN annotated cells and the partition or cluster definition in Monocle 3, cluster definition provides a better resolution and was used to define poorly segregated partitions. A third round of dimensionality reduction and clustering was performed on 27 poorly segregated clusters. Altogether these three iterations produced a total number of 156 cell partitions/clusters. Finally, the CeNGEN annotated cells included in the clusters were used to assign the clusters to a neuronal ID.

### Pairwise comparison of clusters

To compare the similarity of gene expression between our hermaphrodite dataset and the hermaphrodite L4 dataset generated by CeNGEN, we used the 5000 most variable genes in our dataset to calculate the Spearman correlation of the cell cluster using their average gene expression. The results were plotted in a clustered heat map where the complete-linkage clustering method was calculated for rows and columns using euclidean distances.

### Differentially expressed gene analysis

Several metrics were extracted for all genes and all cells, including expression fraction and average expression (table S6-7). Differentially expressed genes between hermaphrodite and male neurons for each cluster was performed using Seurat’s “FindMarkers” function with default parameters on log-normalized dataset. The tresholds were set to include genes detected in 1% of the cells in either of the two populations and that shows at least 0.1-fold difference on average (log-scale base 2) between the two populations. Finally, this step provided log fold changes, percentage of cells where gene is detected for each sex and p value statistics provided by Wilcoxon Rank Sum test (table S3).

### Thresholding of gene expression

In order to define gene expression across neuron types we used the same thresholding strategy as Taylor, S. R. *et al.* (*20*). Specifically, we generated Receiver Operating Characteristic (ROC) curves plotting the True Positive Rate (TPR) against the False Positive Rate (FPR) by comparing our single-cell data to the ground truth. As the ground truth dataset, we used the expression patterns of 160 neuronal genes determined with high confidence in adult hermaphrodite, as previously described(*20*, *45*, *47*, *90–92*). For threshold determination, we used the proportion of cells in which a gene was detected. Genes expressed in at least 1% of cells within each neuron cluster were considered neuronally expressed, whereas those detected in no more than 2% of cells per cluster were considered non-neuronal. For genes that fall between these extremes, we applied percentile thresholding on a gene-by-gene basis, defining thresholds as a fraction of the highest proportion of expressing cells for each gene. Using the R package boot(*93*, *94*), we produced 5,000 stratified bootstraps of the ground truth genes for each threshold percentile. Subsequently, we determined the TPR, FPR, and False Discovery Rate (FDR) for the different thresholds (Fig. 1D-E). For easier comparison, we used the same 4 thresholds of increasing stringency as in (*20*). To define the minimum number of cells required for robust classification performance (area under the ROC curve > 0.8), we generated exploratory ROC curves by progressively including clusters with increasing minimum cell numbers. Even clusters with 4 cells displayed robust classification performance, supporting the reliability of the gene expression tables even for low cell count. At threshold level 2, we noticed that TPR reached 0.8 for cluster containing 9 or more cells. Therefore, only cell types with a minimum of 9 cells both sexes (65 cell types) were kept to define gene expression thresholds. This gives good predictive values with similar performance as reported before for the whole L4 dataset (Fig. 1E). Table S2 contains a list of all the sex-shared clusters obtained with information on the number of neuronal classes they correspond to and the number of cells for each cluster.

### Accuracy of the gene expression table

To evaluate the quality of the gene expression tables at threshold level 2 -used in this work- we assessed their accuracy by comparing with the gene expression atlas for each sex individual neuron type using balanced accuracy, which includes sensitivity (or TPR) and specificity (1-FPR) to correct for dataset imbalance(*95*). Balanced Accuracy = (sensitivity + specificity) / 2. The balanced accuracy increased with the number of cells per cluster (fig. S2D). A one-sided hypothesis test was computed to evaluate whether the overall accuracy rate is greater than the rate of the largest class(*96*) and p-values were adjusted using Holm-Bonferroni correction(*97*). Statistically convincing predictions are achieved for clusters with 9 or more cells.

### Ranking analysis

To rank the sex-shared neuronal clusters according to the number of differentially expressed genes (DEGs) between the sexes, we counted the number of DEGs in each cluster using Python 3.12. Clusters with mixed or unknown identities were removed from this analysis because of the inability to calculate connectivity numbers for these clusters for the analysis below. Clusters were included only if there were more or equal to 9 cells in a cluster for both sexes. This yielded a list of 62 sex-shared clusters we continued analyzing (table S2,5). A gene was considered as differentially expressed if the following criteria were met: average log fold change > 1, p-value < 0.05, threshold ≥ 2 in at least one sex (in order not to exclude genes with no expression in one sex and high expression in the other). Then, the number of DEGs received after filtering (2215) was used to generate gene ontology analysis, calculate the fraction of DEGs per gene family and the number of sex-biased DEGs according to gene family (Fig. 2B-D) and bubble plots representing differential expression of specific gene families (Fig. 3A, Fig. 4A-B, fig. S4, fig. S6) were generated using GraphPad prism 10 or Python 3.12. Genes encoding neuropeptides were defined based on(*41*) and Wormbase (http://www.wormbase.org, release WS294).

### Sex-bias of differentially expressed genes across gene families

Using Python 3.12, we calculated the number of DEGs in males and in herms separately according to gene families (see gene families in Fig. 2D) for all the DEGs between the sexes that passed the filter (Fig. 2A). If a gene appeared in multiple clusters as a DEG, it was counted according to the number of clusters it appeared. For example, *flp-7* is DE in the clusters AVG, RMF and URX, so it was counted three times. For statistical analysis, we performed Fisher’s exact test for each gene family (subtracting the number of each count in each sex from the total count of genes in all families included in the analysis) and then corrected with Bonferroni by multiplying each p-value by the number of comparisons.

### Binary analysis

Using a binary gene expression table for each sex based on threshold 2 stringency (see above, table S6), we counted the number of genes that are either ‘on’ (expressed) in hermaphrodites or ‘off’ (not expressed) in males and vice versa in each of the 62 sex-shared clusters to generate Fig. 2E. Then, this list of genes was compared to the list of DEGs from Fig. 2A to generate Fig. 2F. The total binary genes in each sex, their intersection, and the exclusive genes in each sex were then compared to the total number of genes obtained in our dataset to calculate the percentage shown in Fig. 2G.

### Neurotransmitter identity analysis

To analyze the expression of neurotransmitter genes in our data set, we used Python 3.12, generating expression bubble plots for the two sexes separately (Fig. 4A-B, using tables S7-8, mean expression values of hermaphrodites and males respectively across sex-shared clusters) and for dimorphic expression of neurotransmitter identity genes (Fig. 4C). Neurons with unknown neurotransmitter identity were removed from this analysis including PVM, ASI, AVF, AVH, AVJ, AVK and BDU, leaving 55 clusters for analysis out of the 62 listed in table S5. Then, to calculate the connectivity of the sex-shared neuronal clusters we used Supplementary information 3 from Cook, SJ. *et al.*(*37*). This calculation included only connectivity observed by the electron microscopy (EM) reconstructions(*37*). For each cluster, we calculated using Python 3.12 for each sex the number of outgoing connections (when the neuron is pre-synaptic), the number of incoming connections (when the neuron is post-synaptic) and the sum of these two calculations (all synaptic connections, chemical connections) (table S9). For our analysis we combined left and right connectivity numbers for each neuron. Then, we calculated the difference between the number of male connections and the number of hermaphrodite connections in each cluster, for each synaptic category. These numbers are represented in Fig. 4C as an absolute number. To correlate the difference in connectivity with average log fold change of neurotransmitter identity genes (table S3), we used two-tailed Pearson correlation analysis in GraphPad Prism 10. We did this analysis for every synapse category (outgoing, incoming and all). Clusters with two neurotransmitter identity genes showing dimorphic expression were included twice in the correlation analysis. SIA cluster was discarded because the two neurotransmitter identity genes were showing different sex biases (*cha-1* higher in hermaphrodites and *unc-17* higher in males). AIM was also discarded because its neurotransmitter identity is known to be dimorphic, being glutamatergic in hermaphrodites and cholinergic in males in the adult stage(*35*).

### Microscopy

Confocal images were acquired using Zeiss LSM 880 confocal microscope using 20x, 40x or 63x oil immersion objectives. Worms were anaesthetized in sodium azide (NaN3, final concentration 200 mM, prepared in M9 buffer) and mounted on a 5 % agarose pad (freshly prepared) on a glass slide. The images were processed using ImageJ/Fiji software version 2.3.0/1.53q(*98*) or Zen software. All images of the same group were imaged using identical microscope settings. Figures were prepared using Adobe Illustrator v24.2.1. For imaging NeuroPAL worms, channels were pseudo-colored in accordance with ref(*45*, *46*). For single neuron expression quantification, ROIs were chosen manually at the z-plane with the strongest signal using the respective fluorescent marker. For *sng-1::GFP* analysis we measured the fluorescence intensity of the signal in PLM neurite (observed by *mec-4p::*BFP) measured along 100 µm from the soma. For PLM development analysis we measured the length of the anterior and posterior neurites using *mec-4p::BFP* labeling. For Fig. 6J-K we measured the length of PHC neurite from the anus which is visible through *eat-4p11::GFP*. For fig. S9 we identified AVG based on its anatomic position and shape. For Fig. 7G we calculated the percentage of animals with mislocalized signal of *SNB-1::Venus* and for Fig. 7H we measured *SNB::Venus* signal by manually selecting ROI and normalizing the mean intensity value received to its area for each animal. For Fig. 7I we counted the number of CLA-1 puncta that surround CEP neurons. For RIA axon length analysis, *glr-3p::BFP* images were taken and the most visible RIA was chosen for length measurement using ImageJ/Fiji.

### Neuron identification using NeuroPAL

For neuronal identification the relevant CRISPR GFP reporters were crossed with NeuroPAL landmark strain OH15262 harboring *otIs669* transgene. For imaging NeuroPAL worms, the protocol followed are in accordance with ref(*45*, *46*). For CRISPR GFP neuronal identity, colocalization with the NeuroPAL landmark strain OH15262 harboring *otIs669* transgene was used to determine the identity of all neuronal expression. Using ImageJ version 1.52p, the z-plane with the strongest signal was used to measure the fluorescence intensity of individual neurons that were identified by reporter/stain expression.

### Gene ontology analysis

The WormCat gene set enrichment analysis tool was used for functional annotation of gene ontology terms that are over-represented in differentially expressed genes combined or for each sex(*99*).

### Molecular cloning

To generate *mec-4p::BFP* and *eat-4p11::BFP* for PLM and PHC visualization we used PCR fusion(*100*). *mec-4p* was amplified from *mec-4p::GCaMP3::tagRFP* kindly provided by Doug Kim, while *eat-4p11* was amplified from an eat-4p11 containing plasmid. *BFP* fragment was amplified from a plasmid containing *flp-6::tagBFP* kindly provided by Hannes Bulow. To generate PLM specific rescue of MEC-8, *F10A3.11* promoter (460 bp upstream to *F10A3.11* sequence) amplified from genomic DNA was fused to *mec-8* cDNA amplified from a N2 mixed-stage cDNA library and to *SL2::GFP* amplified from pMO46(*17*) using PCR fusion. *F10A3.11* is predicted by the CeNGEN and our dataset to be specific to PLM in hermaphrodites, and in males it is predicted to be expressed in PLM and RIV. Our observations show that its expression is specific in the tail, but it is expressed in 2-3 additional neurons in the head. *capa-1p* (2154 bp upstream to *capa-1* sequence) and *F32B3.5*p (2805 bp upstream to *F32B4.5* sequence) were amplified from genomic DNA and fused to *GFP::H2B* amplified from genomic DNA of *ins-39::GFP::H2B* knock-in animals(*31*) using PCR fusion. *glr-3p* for generating *glr-3p::BFP* (RIA labeling) was amplified from gDNA. For *dat-1p::novoGFP::cla-1* generation we amplified *dat-1* promoter from a *dat-1p::IRE-1* plasmid(*101*) and *novoGFP::cla-1* from a plasmid kindly provided by Ithai Rabinowitch and Peri Kurshan.

### Touch assays

Gentle touch assays were based on(*102*). In brief, L4 animals were isolated the day before the experiment and stored at 20°C overnight. On the day of the experiment, single 1-day adult animals were transferred into NGM plates freshly seeded with 30 μl OP50. After 30 minutes of habituation, animals were tested by applying a touch to the posterior part of the animal (anteriorly to the anus) with an eyelash. Touch was applied when worms were not moving or moved very slowly. Each worm was tested five times with intervals of at least 10 seconds between each trial. The tested worm was given a score of one if it moved forward in response to the touch, and zero if it did not move forward. The response index was then calculated as the average of the forward responses. For anterior gentle touch responses, the anterior part of the animal was touched with an eyelash and a reverse movement response was scored. For harsh tail touch assays, animals were tested by applying a touch to the tail with a flattened platinum wire pick as previously described(*103*) and their forward movements were scored similarly to animals subjected to posterior gentle touch.

### Single Molecule Fluorescent in Situ Hybridization

*mec-8* CAL Fluor Red 590 nm custom was designed using Stellaris Designer. All the reagents were ordered from Biosearch Technologies and most of the preparations are based on their protocol (Stellaris RNA FISH – protocol for *C. elegans*), with minor adjustments. Synchronized L4 worm population expressing *mec-4p::GFP (zdIs5)* was washed three times with M9 to remove bacterial residues. Then, after aspirating the M9, 1 mL of fixation buffer (4% formaldehyde in 1XPBS) was added to the worms followed by 45 minutes incubation at room temperature. After fixation, worms were washed twice with 1XPBS and supplemented with 70% ethanol. Worms were stored at 4°C overnight. The next day, larvae pellet was centrifuged, and the ethanol was discarded. Worms were then added with 1 mL Wash Buffer A (2 mL Stellaris RNA FISH Wash Buffer A (Biosearch Technologies Cat# SMF-WA1-60), 7 mL Nuclease-free water, 1 mL Deionized Formamide) and incubated at room temperature for 5 minutes. After discarding Wash Buffer A (spin down), worms were dispensed with 100 μL of the Hybridization Buffer containing probe (2 µL of probe set (12.5 µM in TE buffer)) into a mixture of 100 µL Hybridization Buffer (450 µL Stellaris RNA FISH Hybridization Buffer, Biosearch Technologies Cat# SMF-HB1-10 and 50 µL Deionized Formamide) to achieve 250 nM probe solution and incubated at 37°C in the dark overnight. In the morning, worms were washed once with Wash Buffer A and incubated with Wash Buffer A at 37°C in the dark for 30 minutes. Then, Wash Buffer A was discarded, and worms were added 1 mL DAPI (4’,6-diamidino-2-phenylindole, Wash Buffer A consisting of 5 ng/mL DAPI) to stain the nuclei and incubated at 37°C in the dark for 30 minutes. Then, DAPI was discarded, and worms were added with 1 mL of Wash Buffer B (Biosearch Technologies Cat# SMF-WB1-20) and incubated for 5 minutes at room temperature. After discarding Wash Buffer B, 30 µL Vectashield® Mounting Medium (Vector Laboratories Cat #H-1000) were added to the worms for imaging. Imaging was done using Zeiss LSM 880 confocal microscope with 63x oil immersion objective. To image large sample size, we stored samples at 4°C for a few weeks with no signal loss. To differentiate between hermaphrodites and male at all larval stages, we used *ins-39*::SL2::GFP::H2B CRISPR knock-in worms as described previously(*31*). For quantification, we followed a protocol based on(*104*). We performed Gauss blur filter of sigma = 1 for every image. Then, we performed for each image “Max Entropy” thresholding in Fiji that was selected based on trial-and-error process done on 2-3 representative images. Then, we carried particle analysis for the image in the ROI determined by the GFP channel. This gives a count for the number of particles in a specific ROI. We also performed manual counting of smFISH puncta and got the same results.

### Calcium imaging

To image spontaneous calcium traces in PLM, we used animals expressing *mec-4p::GCaMP3.35*. Using a drop of S-basal with 10 mM levamisole that was filtered with 0.22 um filter, adult worms were inserted in a dual olfactory chip that was a kind gift from Michael Krieg lab, fabricated as previously described(*105*, *106*). The flow rate in the chip was 0.005 ml/min. Imaging was done with a Zeiss LSM 880 confocal microscope using a 40x magnification water objective. When the worm was properly located inside the chip, PLM was imaged for 1150 frames (2.8 minutes), with imaging rate of 6.667 Hz. For analysis, the GCaMP fluorescence intensity was measured using FIJI. ROIs of PLM soma were marked manually to extract mean intensity values. Data analysis was performed using Python 3.12. For each worm, the baseline fluorescent level (F0) was calculated by averaging the mean values of 100 frames in the beginning of each recording. Then, for each frame, ΔF was calculated by subtracting F0 from the value of that time point, and the result was divided by F0, to normalize the differences in the fluorescence baseline levels between individuals (ΔF/F0). To calculate the differences between the sexes in PLM basal activity, we performed Fourier transform on the ΔF/F0 values, then the amplitudes were normalized for each worm to its own maximum value. Finally, AUC (area under the curve) was measured using the trapz function in Python, using frequencies up to 2 Hz.

### RNAi-Induced Gene Silencing

Gene silencing using RNA interference was carried out as described previously using the feeding protocol on agar plates(*107*). Hermaphrodites at L4 stage were fed on HT115 bacteria expressing dsRNA (source Ahringer library) for the targeted genes or on control bacteria harboring empty RNAi vector, L4440. Posterior (gentle) touch responses at 1-day adult from their progeny were scored. The RNAi efficiency was measured using *pos-1* RNAi as a positive control. For every RNAi experiment, a fresh *pos-1* control was prepared.

### Optogenetic stimulation assay

To measure the response of mechanosensory stimulus to the head and tail of hermaphrodite and male *C. elegans* worms, we used AML470(*108*) strain that expresses light-gated ion channel Chrimson and a fluorescent reporter mCherry in all the six gentle touch neurons under the control of *mec-4* promoter. L4 males and hermaphrodites were isolated a day prior to the day of experiments. Males were separated from hermaphrodites to make sure that they are not mated. The worms were transferred to freshly seeded plates consisting of 1 ml of 0.5 mM all-trans-retinal (Sigma-Aldrich Cat. # R2500) mixed with *E. Coli* (OP50) as food source and stored in the dark at 20°C until the next day. Day 1 adult male and hermaphrodite worms were used in the optogenetic behavior assay. On the day of experiment 20-30 worms were transferred to the unseeded agarose plates to carry out behavior assay while delivering optogenetic stimulation. To measure the response of mechanosensory stimulus to head and tail of male and hermaphrodites, we delivered optogenetic stimulus to the worms using a high throughput optogenetic delivery system(*108*, *109*). Optogenetic stimulation protocol was same as described in the previous work(*108*). Briefly, a 400-µm diameter red light stimulation was delivered to either the head, tail, or both simultaneously to each tracked worm on the agarose plate. The illumination spot was centered on the tip of the head or tail of the worm. Optogenetic stimulus of 630 nm wavelength was delivered through a computer-controlled projector for 1 second with an inter trigger stimulus interval of 30 seconds. The stimulus intensity was randomly drawn independently for the head and the tail from the set of 0 and 80 µW/mm^2^ intensities. Thus, there were four stimulus combinations: 1) Head: 0 µW/mm^2^, Tail 0 µW/mm^2^ 2) Head: 0 µW/mm^2^, Tail 80 µW/mm^2^, 3) Head: 80 µW/mm^2^, Tail 0 µW/mm^2^, and 4) Head: 80 µW/mm^2^, Tail 80 µW/mm^2^. Each behavior assay lasted for 30 minutes. The analysis of camera frames of the behavior assay was same as described in the previous studies(*108*, *109*). Briefly, two independent sets of behavior mapping algorithms were employed, one for real-time tracking of worms and optogenetic stimulus delivery, and the other for post processing analysis. The real-time algorithm in LabVIEW tracked all the worms on the agarose plate and calculated various behavior parameters of each worm such as velocity, body curvature, centerline, etc. The first and last index of the centerline were assigned as head and tail respectively. Every 30 seconds, the real-time algorithm signals the computer-controlled projector to deliver optogenetic stimulus to the head, tail, or both simultaneously for every tracked worm with one of the four stimulus intensity combinations. In the post-processing algorithm, we correct for any error in the assignment of head and tail of the worm. This algorithm also classifies behavior of the worm by classifying pose dynamics into a behavior map of forward, reverse, and turns(*109*, *110*). Same analysis steps were used as in the previous study to determine the change in velocity(*108*). The velocity of the worm two seconds prior to stimulus onset was defined as the initial velocity of the worm. The velocity of the worm at the end of one second stimulus was defined as the final velocity. Change in velocity for each stimulus event for a worm was defined as the difference between the final velocity and the initial velocity.

### Correlation analysis

The entire analysis was done using Python 3.12. Using the number of outgoing connections calculated for Fig. 4 (See table S9), we generated for each sex a matrix including the number of outgoing connections and mean expression values of all genes taken from table S7-8 across the 62 sex-shared neuronal clusters (table S5). Genes that were not expressed in any of the 62 clusters were removed from the analysis. Then, we performed Pearson correlation calculation with Benjamini-Hochberg False Discovery Rate (FDR) correction for each comparison to account for multiple comparisons, providing correlation coefficient for each gene in the data set of hermaphrodites and male separately. We refer in the text to significantly correlated genes to genes with p-valued adjusted < 0.05.

## Acknowledgments

We thank members of the Oren-Suissa lab and Laurent lab for their critical insights regarding the manuscript. We thank Dr. Joseph Dicken and Dr. Efrat Hagai at Flow Cytometry Unit, Life Science Core Facility, Weizmann Institute of Science for helping with FACS sorting. scRNA-seq library preparation was done with critical advice from Dr. Hadas Keren-Shaul, Dr. Merav Kedmi and Dr. David Pilzer from the Genomics Sandbox unit at the Life Science Core Facility of Weizmann Institute of Science. We thank Sapir Sela from the Oren-Suissa lab for cloning the *dat-1p::novogfp::cla-1* plasmid. We are grateful to Oliver Hobert lab for sharing strains. Some strains used in this study were obtained from *Caenorhabditis* Genetics Center (CGC), which is funded by the NIH Office of Research Infrastructure Programs (P40 OD010440). We thank WormAtlas which is supported by NIH OD 010943. We are thankful to Doug Kim, Ithai Rabinowitch, Peri Kurshan, Hannes Buelow labs for providing plasmids. We thank Michael Krieg lab for providing the dual-olfactory chips. We thank WormBase, an online biological database for *C. elegans*, which is supported by Grant U41 HG002223 from the National Human Genome Research Institute at the NIH, the UK Medical Research Council, and the UK Biotechnology and Biological Sciences Research Council. Figures 1A, 1F, 5A, 5I, and 6A, were created with BioRender.com.

## Author contributions

Conceptualization: R.H., H.S., Y.S., G.G., E.S., P.L., M.O.S. Formal analysis: R.H., H.S., R.L., R.R., G.S., S.K., P.L., M.O.S. Methodology: R.H., H.S., R.L., Y.S., G.S., P.L., M.O.S. Investigation: R.H., H.S., Y.S., S.N.H., S.K., P.L., M.O.S. Project administration: R.H., H.S., M.O.S. Software: R.L., G.S., R.R. Resources: R.H., H.S., G.S., S.N.H., P.L., M.O.S. Supervision: A.L., E.S., P.L., M.O.S. Validation: R.H., H.S., Y.S., S.N.H., A.L., P.L., M.O.S. Visualization: R.H., H.S., R.L., G.S., P.L., M.O.S. Writing – original draft: R.H., H.S., R.L., Y.S., P.L., M.O.S. Writing – review and editing: R.H., H.S., R.L., Y.S., G.S., R.R., G.G., S.K., A.L., P.L., M.O.S. R.H., H.S., and R.L are co-first authors.

## Data and Materials Availability

All data needed to evaluate the conclusions in the paper are present in the paper and/or the Supplementary Materials. The link to codes used in this paper is available in the following repository: https://doi.org/10.34933/60314fe3-7cbc-4aaf-a22c-f29799b7a137. The sequence data of this article have been submitted to NCBI and are available under GEO accession number GSE286418 (https://www.ncbi.nlm.nih.gov/geo/query/acc.cgi?acc=GSE286418).

