## Supplemental Information for "Decoding sexual dimorphism of the sex-shared nervous system at single-neuron resolution"

Rizwanul Haque *et al.*

**This PDF file includes:**

Supplementary table of contents
Figs. S1 to S10
Supplementary Tables S1-S10

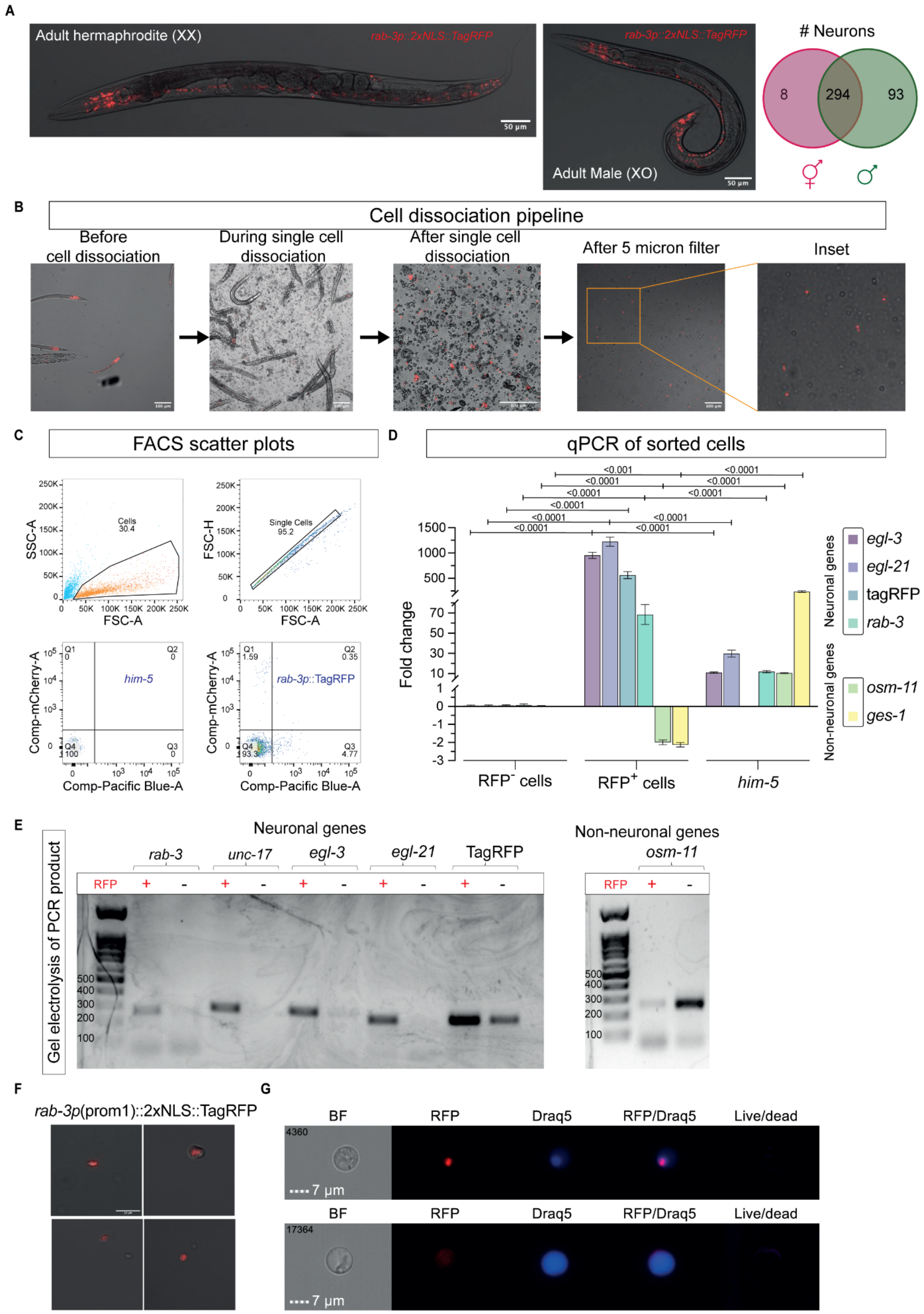
A
Adult hermaphrodite (XX)
rab-3p::2xNLS::TagRFP
50 μm
Adult Male (XO)
### Neurons
8
294
93
B
Cell dissociation pipeline
Before cell dissociation
During single cell dissociation
After single cell dissociation
After 5 micron filter
Inset
C
FACS scatter plots
SSC-A
FSC-A
Cells 30.4
FSC-H
Single Cells 95.2
Comp-mCherry-A
Comp-Pacific Blue-A
him-5
rab-3p::TagRFP
Q1 0
Q2 0
Q3 0
Q4 100
Q1 1.59
Q2 0.35
Q3 4.77
Q4 93.3
D
qPCR of sorted cells
Fold change
<0.001
<0.0001
RFP⁻ cells
RFP⁺ cells
him-5
Neuronal genes
egl-3
egl-21
tagRFP
rab-3
Non-neuronal genes
osm-11
ges-1
E
Gel electrolysis of PCR product
Neuronal genes
rab-3
unc-17
egl-3
egl-21
TagRFP
RFP
+
-
500
400
300
200
100
Non-neuronal genes
osm-11
F
rab-3p(prom1)::2xNLS::TagRFP
G
BF
RFP
Draq5
RFP/Draq5
Live/dead
4360
17364
7 μm

**Fig. S1. Single-cell quality assessment Protocol**

(A) Confocal images of the pan-neuronal marker, *rab-3p(prom1)::2xNLS::TagRFP* in the two adult sexes of *C. elegans*. The pan neuronal marker labels all sex-shared neurons in both sexes. The Venn diagram compares the number of neurons that are shared or sex-specific. 294 neurons are present in both sexes, corresponding to 116 molecular classes. Scale bars represent 50 µm. (B) Single cell dissociation process of worms expressing the pan-neuronal marker, *rab-3p(prom1)::2xNLS::TagRFP*. The figure displays representative confocal micrographs of before, during, and after the dissociation process, as well as following passage through a 5-micron filter. Scale bars represent 100 µm. (C) Scatterplot depicting the FACS profile for cells dissociated from the strains expressing pan-neuronal *TagRFP* marker and *him-5* reference strain. eFlour 450 dye was used to discriminate live and dead cells. Gating was conducted based on side scatter (SSC)-Area vs forward scatter (FSC)-Area, followed by the selection of single cells by discriminating against doublets using FSC-Height vs FSC-Area. Wild-type *him-5* cells without RFP+ expression served as a control for gating. (D) qPCR analysis of neuronal (*egl-3*, *egl-21*, tagRFP and *rab-3*) and non-neuronal (*osm-11* and *ges-1*) genes/markers in equal amount of cells sorted based on the presence (RFP+) or absence (RFP-) of TagRFP. (E) Gel electrophoresis of PCR products amplified using primers for neuronal (*rab-3*, *unc-17*, *egl-3*, *egl-21* and TagRFP) and non-neuronal (*osm-11*) cDNAs isolated from equal amounts of RFP+ and RFP- sorted cells. Neuronal mRNA is highly enriched in RFP+ cells and non-neuronal mRNA is highly enriched in RFP- cells. (E) Representative confocal micrographs of RFP+ cells after FACS sorting. Scale bars represent 10 µm. (F) Representative micrographs of RFP+ cells (top) RFP- cells (bottom) after sorting using ImageStream Flow Cytometer. Scale bars represent 7 µm.

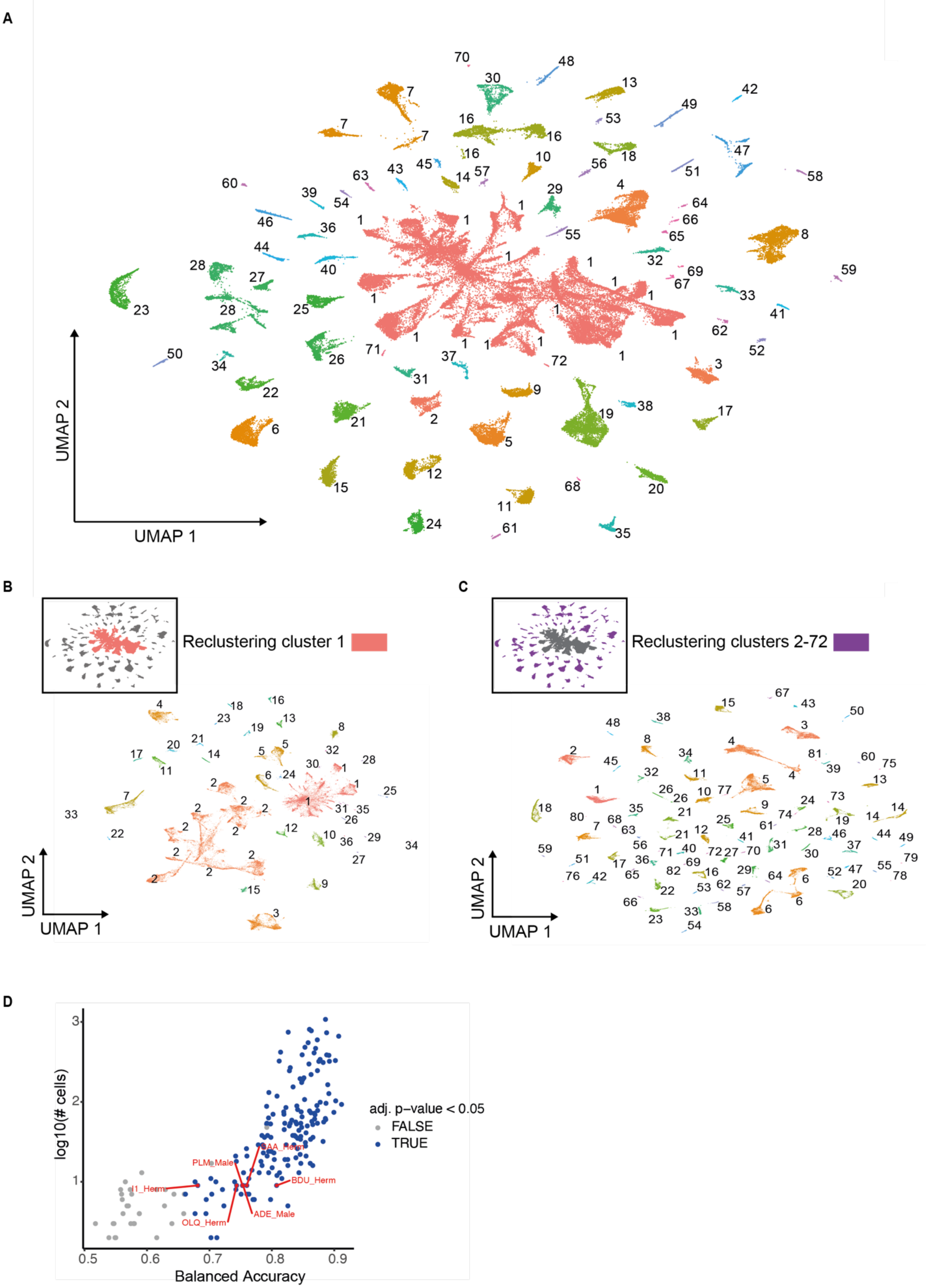
A
UMAP 1
UMAP 2
B
Reclustering cluster 1
UMAP 1
UMAP 2
C
Reclustering clusters 2-72
UMAP 1
UMAP 2
D
log10(# cells)
Balanced Accuracy
adj. p−value < 0.05
FALSE
TRUE
I1_Herm
PLM_Male
SAA_Herm
BDU_Herm
OLQ_Herm
ADE_Male

**Fig. S2. Iterative clustering analysis of *C. elegans* single cell populations using UMAP**
(A) UMAP representation of 72 cells clusters identified in round 1 of clustering (Monocle parameters in methods). Numbers represent individual cluster. (B) UMAP representation of 36 cells clusters after re-clustering of cluster 1 (Monocle parameters in methods). Inset shows the cluster/s that underwent round 2 of clustering. (C) UMAP representation of 82 cells clusters after re-clustering of cluster 2-72 (Monocle parameters in methods). Inset shows the cluster/s that underwent round 2 of clustering. (D) Balanced accuracy using threshold 2 for individual neurons vs cell. For each individual neuron type, the balanced accuracy value is plotted against the number of cells in the associated cluster. Significant accuracies are highlighted in blue for each neuron type. Clusters containing 9 cells are highlighted in red.

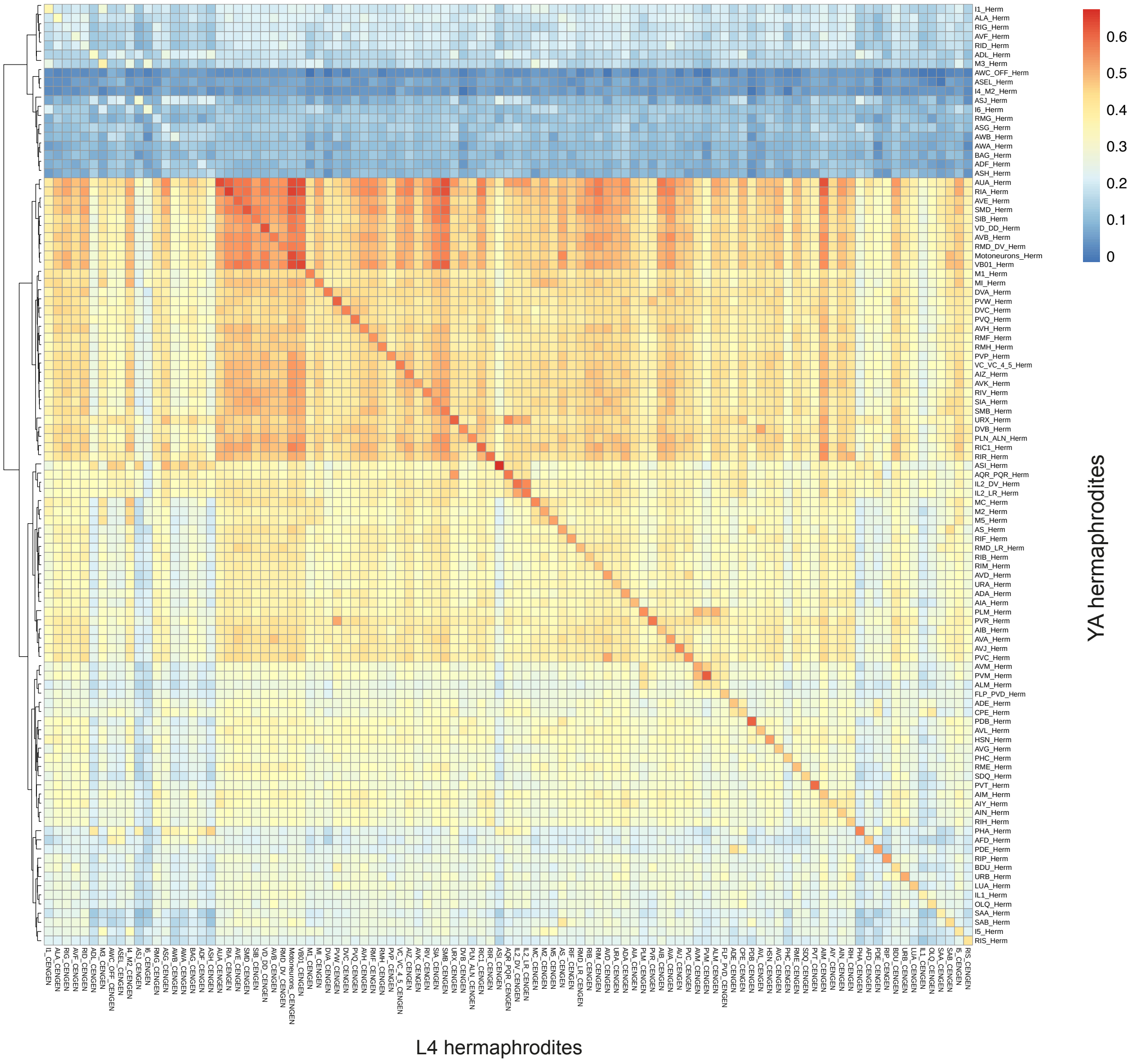


**Fig. S3. Comparative heatmap reveals high similarity index of YA and L4 hermaphrodite clusters**

Clustered heatmap by Spearman correlation using the mean expression of highly variable genes shows relationship between clusters from YA hermaphrodites (current datasets) and previously identified L4 hermaphrodites clusters from CeNGEN(*20*). There is a high similarity index between known clusters from both the datasets. Hierarchical clustering of rows (Hermaphrodite dataset) was performed using Euclidean distances and the complete linkage method. Columns (CeNGEN dataset) were sorted according to the order of rows.

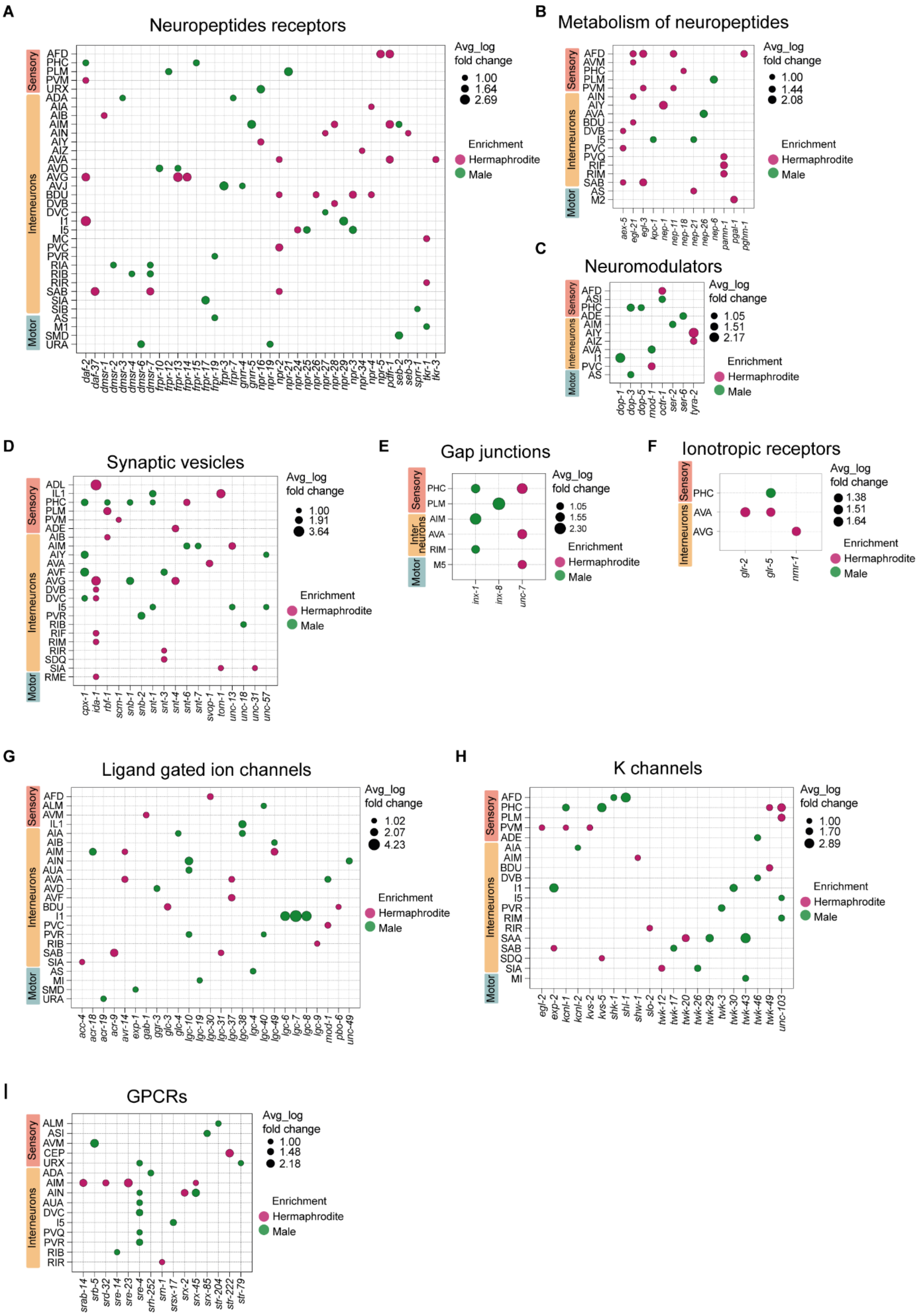
A
Neuropeptides receptors
Avg_log fold change
1.00
1.64
2.69
Enrichment
Hermaphrodite
Male
Sensory
Interneurons
Motor
AFD PHC PLM PVM URX ADA AIA AIB AIM AIN AIY AIZ AVA AVD AVG AVJ BDU DVB DVC I1 I5 MC PVC PVR RIA RIB RIR SAB SIA SIB AS M1 SMD URA
daf-2 daf-37 dmsr-1 dmsr-2 dmsr-3 dmsr-4 dmsr-6 dmsr-7 frpr-10 frpr-12 frpr-13 frpr-14 frpr-15 frpr-17 frpr-19 frpr-3 frpr-7 gnrr-4 gnrr-5 npr-16 npr-19 npr-2 npr-21 npr-24 npr-25 npr-26 npr-27 npr-28 npr-29 npr-3 npr-34 npr-4 npr-5 pdfr-1 seb-2 seb-3 sprr-1 tkr-1 tkr-3
B
Metabolism of neuropeptides
Avg_log fold change
1.00
1.44
2.08
Enrichment
Hermaphrodite
Male
Sensory
Interneurons
Motor
AFD AVM PHC PLM PVM AIN AIY AVA BDU DVB I5 PVC PVQ RIF RIM SAB AS M2
aex-5 egl-21 egl-3 kpc-1 nep-1 nep-11 nep-18 nep-21 nep-26 nep-6 pamn-1 pgal-1 pghm-1
C
Neuromodulators
Avg_log fold change
1.05
1.51
2.17
Enrichment
Hermaphrodite
Male
Sensory
Interneurons
Motor
AFD ASI PHC ADE AIM AIY AIZ AVA I1 PVC AS
dop-1 dop-3 dop-5 mod-1 octr-1 ser-2 ser-6 tyra-2
D
Synaptic vesicles
Avg_log fold change
1.00
1.91
3.64
Enrichment
Hermaphrodite
Male
Sensory
Interneurons
Motor
ADL IL1 PHC PLM PVM ADE AIB AIM AIY AVA AVF AVG DVB DVC I5 PVR RIB RIF RIM RIR SDQ SIA RME
cpx-1 ida-1 rbf-1 scm-1 snb-1 snb-2 snt-1 snt-3 snt-4 snt-6 snt-7 svop-1 tom-1 unc-13 unc-18 unc-31 unc-57
E
Gap junctions
Avg_log fold change
1.05
1.55
2.30
Enrichment
Hermaphrodite
Male
Sensory
Inter neurons
Motor
PHC PLM AIM AVA RIM M5
inx-1 inx-8 unc-7
F
Ionotropic receptors
Avg_log fold change
1.38
1.51
1.64
Enrichment
Hermaphrodite
Male
Sensory
Interneurons
PHC AVA AVG
glr-2 glr-5 nmr-1
G
Ligand gated ion channels
Avg_log fold change
1.02
2.07
4.23
Enrichment
Hermaphrodite
Male
Sensory
Interneurons
Motor
AFD ALM AVM IL1 AIA AIB AIM AIN AUA AVA AVD AVF BDU I1 PVC PVR RIB SAB SIA AS MI SMD URA
acc-4 acr-18 acr-19 acr-9 avr-14 exp-1 gab-1 ggr-3 glc-3 glc-4 lgc-10 lgc-19 lgc-30 lgc-31 lgc-37 lgc-38 lgc-4 lgc-40 lgc-49 lgc-6 lgc-7 lgc-8 lgc-9 mod-1 pbo-6 unc-49
H
K channels
Avg_log fold change
1.00
1.70
2.89
Enrichment
Hermaphrodite
Male
Sensory
Interneurons
Motor
AFD PHC PLM PVM ADE AIA AIM BDU DVB I1 I5 PVR RIM RIR SAA SAB SDQ SIA MI
egl-2 exp-2 kcnl-1 kcnl-2 kvs-2 kvs-5 shk-1 shl-1 shw-1 slo-2 twk-12 twk-17 twk-20 twk-26 twk-29 twk-3 twk-30 twk-43 twk-46 twk-49 unc-103
I
GPCRs
Avg_log fold change
1.00
1.48
2.18
Enrichment
Hermaphrodite
Male
Sensory
Interneurons
ALM ASI AVM CEP URX ADA AIM AIN AUA DVC I5 PVQ PVR RIB RIR
srab-14 srb-5 srd-32 sre-14 sre-23 sre-4 srh-252 srn-1 srsx-17 srx-2 srx-45 srx-85 str-204 str-222 str-79

**Fig. S4. Sexually dimorphic regulation of neuronal gene families**

(A-I) Bubble plot illustration of differentially expressed genes between the two sexes of Neuropeptides receptors (A), Metabolism of neuropeptides (B), Neuromodulators (C), Synaptic vesicles (D), Gap junctions (E), Ionotropic receptors (F), Ligand gated ion channels (G), K channels (H), and GPCRs (I) gene families. Bubble size corresponds to the average fold change in gene expression (only genes meeting the criteria of average log fold change > 1, number of cells in a cluster ≥ 9, p-value < 0.05, threshold ≥ 2 in at least one sex), and bubble color indicates enrichment in either sex (green for male enrichment, pink for hermaphrodite enrichment). Rows represent neuron types, grouped by sensory neurons, interneurons, and motor neurons. Columns represent individual genes.

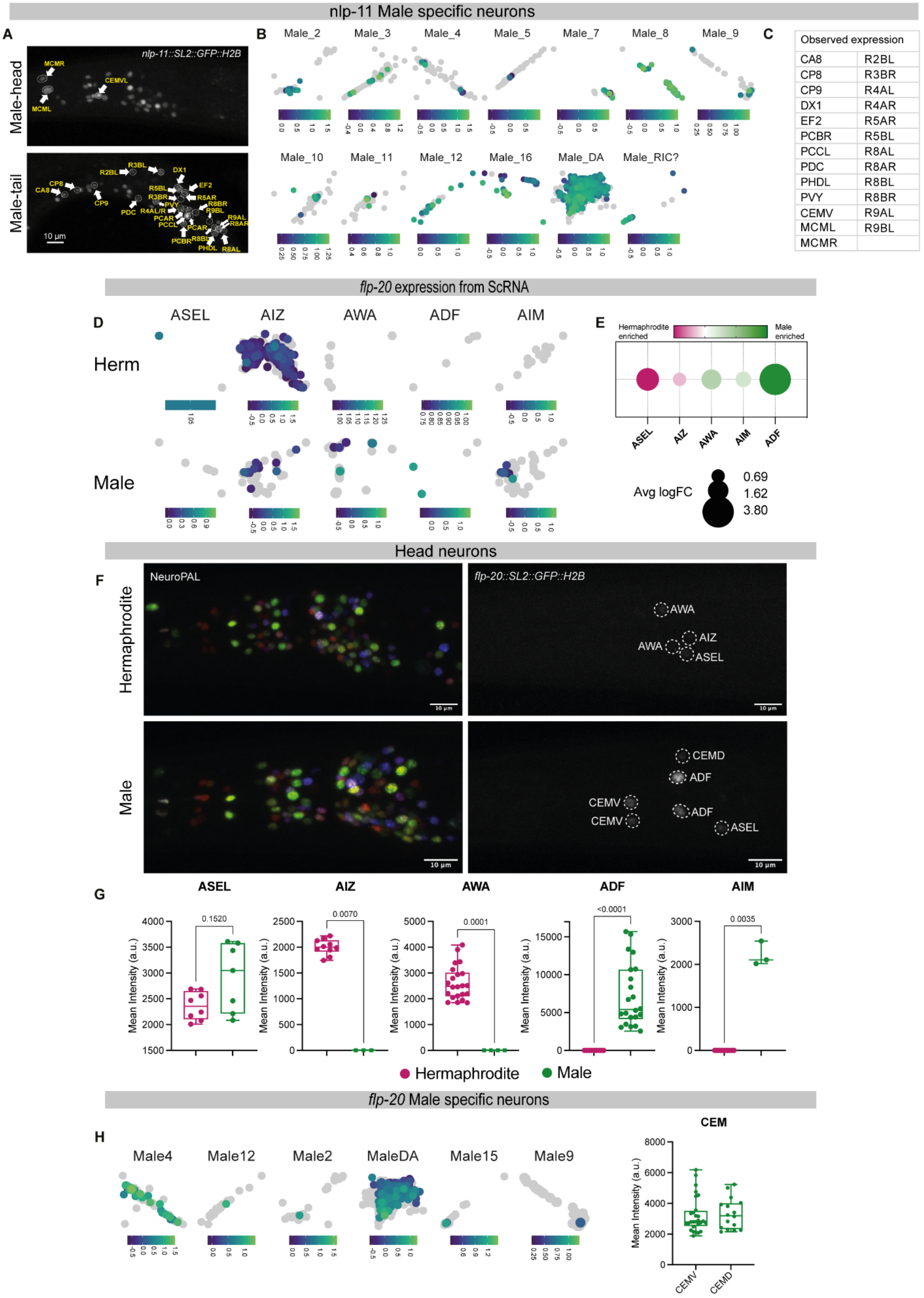
nlp-11 Male specific neurons
A
B
C
Male-head
Male-tail
nlp-11::SL2::GFP::H2B
MCMR
CEMVL
MCML
R3BL
R2BL
DX1
CP8
CA8
CP9
EF2
R5BL
R3BR
R5AR
PVY
R8BR
PDC
R4AL/R
R9BL
PCAR
PCCL
R9AL
R8AR
PCBR
R8BL
PHDL
R8AL
10 µm
Male_2
Male_3
Male_4
Male_5
Male_7
Male_8
Male_9
Male_10
Male_11
Male_12
Male_16
Male_DA
Male_RIC?
Observed expression
CA8 R2BL
CP8 R3BR
CP9 R4AL
DX1 R4AR
EF2 R5AR
PCBR R5BL
PCCL R8AL
PDC R8AR
PHDL R8BL
PVY R8BR
CEMV R9AL
MCML R9BL
MCMR
flp-20 expression from ScRNA
D
E
ASEL
AIZ
AWA
ADF
AIM
Herm
Male
Hermaphrodite enriched
Male enriched
Avg logFC
0.69
1.62
3.80
Head neurons
F
NeuroPAL
flp-20::SL2::GFP::H2B
Hermaphrodite
Male
AWA
AIZ
ASEL
CEMD
ADF
CEMV
10 µm
G
Mean Intensity (a.u.)
0.1520
0.0070
0.0001
<0.0001
0.0035
Hermaphrodite
Male
flp-20 Male specific neurons
H
Male4
Male12
Male2
MaleDA
Male15
Male9
CEM
CEMV
CEMD

**Fig. S5. Neuropeptide validations using NeuroPAL**

(A) UMAP projection of male specific clusters showing expression of *nlp-11* gene in males. Heatmap represent the expression levels. (B) Identified expression of *nlp-11*(*syb4759*[*nlp-11::SL2::GFP::H2B*]) fluorescence intensity from Figure 3E in male-specific clusters. (C) UMAP projection of ASEL, AIZ, AWA, ADF and AIM clusters showing expression of *flp-20* gene in both sexes. Heatmap represent the expression levels. (D) Bubble plot representation of *flp-20* genes expression in ASEL, AIZ, AWA, ADF and AIM clusters of the two sexes. Bubble size represents Avg log fold change, bubble color (green: males and pink: hermaphrodites) represents enrichment in either sex. (E) Representative confocal micrographs showing the distinctive multi-colored neuronal nuclei in the NeuroPAL worm strain otIs669 [NeuroPAL] (left), a strain in which every neuron class can be uniquely identified based on its colour-based barcode(*45, 46*). These images were used to identify neurons expressing the *flp-20*(*syb4049*[*flp-20::SL2::GFP::H2B*]) reporter in young adult hermaphrodites (right, top) and males (right, bottom). Scale bars represent 10μm. (F) Quantification of *flp-20*(*syb4049*[*flp-20::SL2::GFP::H2B*]) fluorescence intensity from E in head neurons ASEL, AIZ, AWA, ADF and AIM in both sexes at YA stage. (G) UMAP projection of male specific clusters showing expression of *flp-20* gene in males. Heatmap represent the expression levels (left). Quantification of *flp-20*(*syb4049*[*flp-20::SL2::GFP::H2B*]) fluorescence intensity from E in male-specific clusters CEMV and CEMD (right). a.u., arbitrary units. Vertical bars in the box-and-whiskers graph represent the median, with dots showing all points from min to max. 16-28 animals were imaged for each group, but some neurons could be identified in a subset of the animals (see below). For each neurons group comparison, n represent the number of animals in each group in which neurons were identified and recorded. n = 8 hermaphrodite, 7 males for ASEL, n = 10 hermaphrodite, 3 males for AIZ, n = 22 hermaphrodite, 4 males for AWA, n = 9 hermaphrodite, 22 males for ADF, n = 10 hermaphrodite, 3 males for AIM. n = 17 animals for CEMD and n = 28 animals for CEMV. a.u., arbitrary units. Scale bars represent 10 μm. In the box-and-whiskers graph, the center line in the box denotes the median, while the box contains the 25th to 75th percentiles of the dataset, whiskers define the minimum and maximum value with dots showing all points. We performed a two-sided Mann-Whitney test for each comparison.

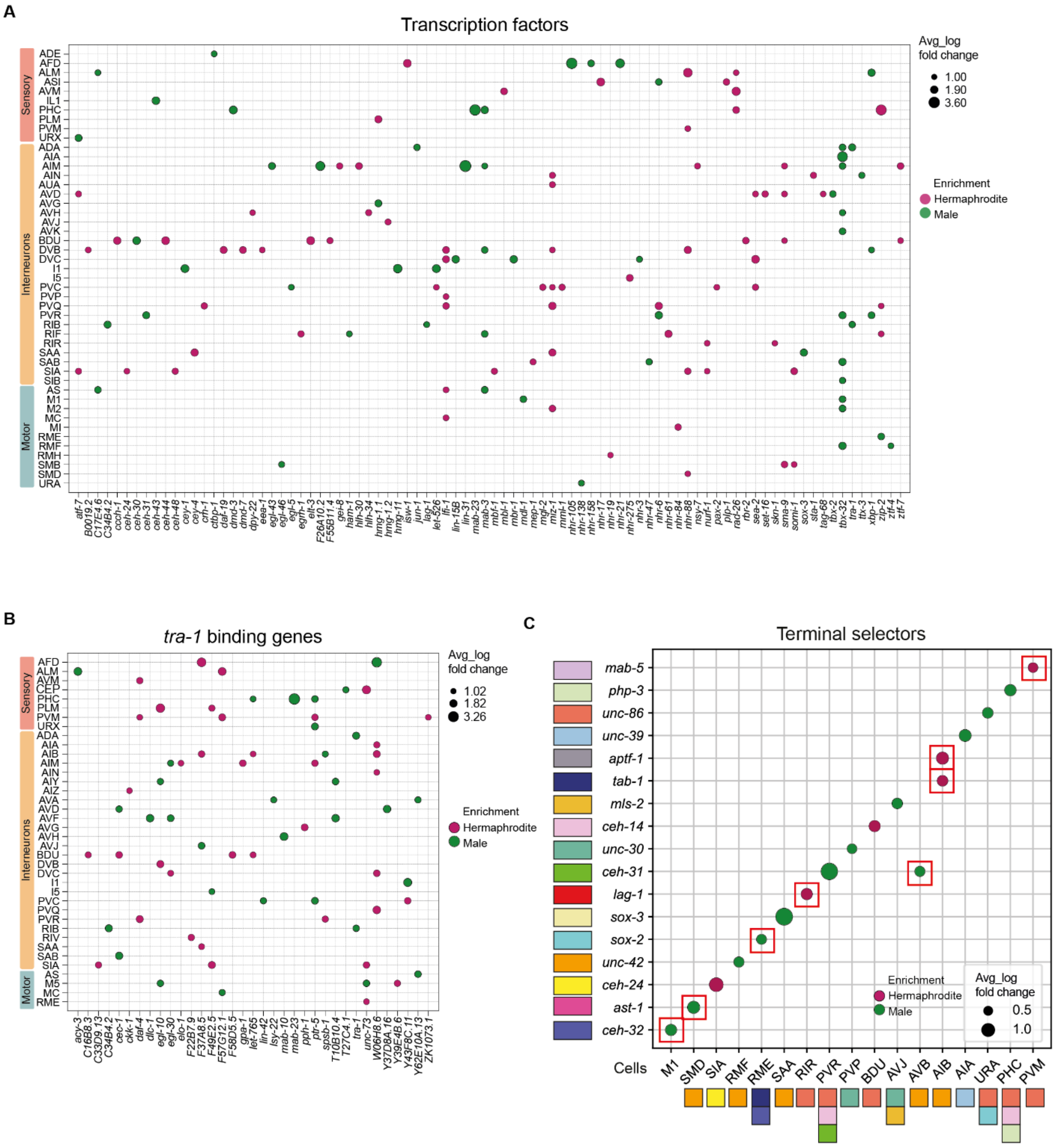


**Fig. S6. Sex-based differential gene expression in transcription factor and related gene families**

(A-C) Bubble plot illustration of differentially expressed genes between the two sexes of transcription factors (A), *tra-1* binding genes (B), and Terminal selectors (C) gene families. In A TFs known to drive sex-specific maturation, such as *tra-1* and *mab-3*, were differentially expressed in only 2 and 4 neurons, respectively. Bubble size corresponds to the average fold change in gene expression. For A-B only genes meeting the criteria of average log fold change > 1, number of cells in a cluster ≥ 9, p-value < 0.05, threshold ≥ 2 in at least on sex. For C only genes meeting the criteria of average log fold change > 0.5, number of cells in a cluster ≥ 9, p-value < 0.05 and fraction of cells expressing the gene in a cluster > 0.5 in at least one sex. Bubble color indicates enrichment in either sex (green for male enrichment, pink for hermaphrodite enrichment). In A-B rows represent neuron types, grouped by sensory neurons, interneurons, and motor neurons and columns represent individual genes. In C rows represent terminal selector genes and columns represent cell types. In C, colors indicate the prediction for terminal selector identity for each cell type: each gene is represented with a different color, and each cell type is represented with one or more colors to indicate the terminal selectors that are required for the identity of this cell type based on literature (*47*, *48*). Red squares represent enrichment of a terminal selector in an unpredicted cell type. Only differentially expressed terminal selectors are shown. Thus, some cell types are missing terminal selectors, such as M1, whose terminal selector is *ceh-34(47)*.

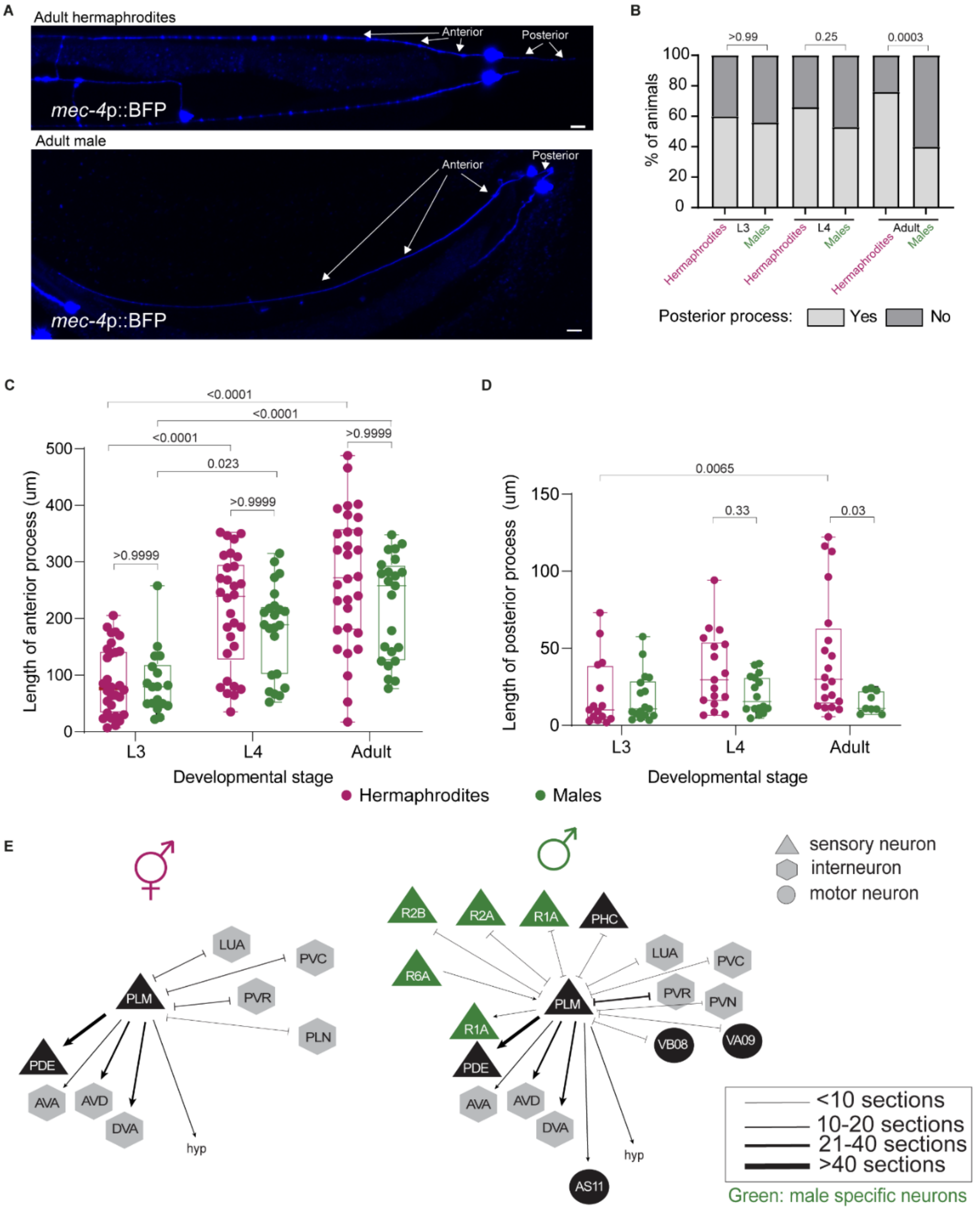


**Fig. S7. Sexually dimorphic morphology of PLM posterior process appears during the adult stage**

(A) Representative confocal micrographs of PLM neurons in adult hermaphrodites and males, using *mec-4p::BFP* labeling. (B) Quantification from A, showing the percentage of animals in each sex with or without posterior PLM processes, during L3, L4 and adult developmental stages. Quantification of the length of PLM anterior process (C) or posterior process (D) in each sex, during L3, L4 and adult developmental stages. (E) Schematic diagram of the connectivity of the PLM neuron in both sexes in the adult stage based on electron microscopy reconstructions (*37*)

Chemical and electrical synapses between sensory (triangles), motor (circles) and inter- (hexagons) neurons are depicted as arrows and inhibitory arrows, accordingly. Arrow thickness correlates with the degree of connectivity (number of sections over which en passant synapses are observed). For (B) we performed two-sided Fisher's exact test followed by p-value Bonferroni correction for multiple comparisons and for (C) and (D) we performed a Kruskal-Wallis test followed by a Dunn's multiple comparison test. In the box-and-whiskers graph (C-D), the center line in the box denotes the median, while the box contains the 25th to 75th percentiles of the dataset, whiskers define the minimum and maximum value with dots showing all points.

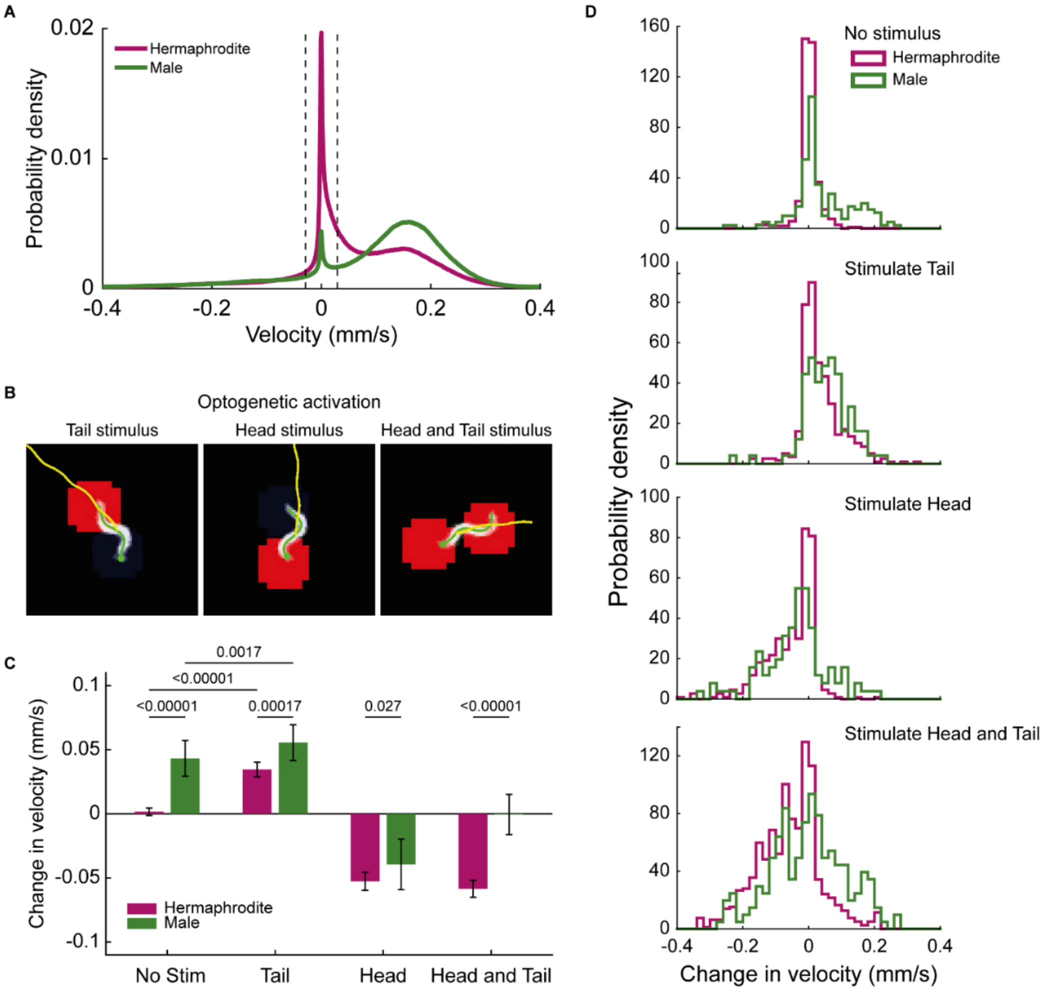


**Fig. S8. Hermaphrodites and males respond differently to optogenetic stimulation of touch receptor neurons**

Hermaphrodites most often respond to combined head and tail stimuli as if they had received a head stimulus, while the response of males to combined head and tail stimulation is more variable: sometimes they respond as if they were stimulated on the head, sometimes they respond as if they were stimulated on the tail. A) Baseline velocity distributions for spontaneous locomotion. Dashed lines denote pre-stim inclusion criteria for subsequent experiments. B) 1s duration optogenetic illumination is delivered to the tail, head or both of animals expressing Chrimson under a *mec-4* promotor. Only those stimuli events that were delivered to animals that were paused or nearly paused are considered (+/-0.029 mm/s, dashed lines from panel A). Computer vision extracted features are shown: head (green circle), body centerline (green line), and locomotion trajectory (yellow line). Optogenetic illumination is delivered in red areas. C) Change in velocity is defined as change in velocity from a pre stimulus baseline (see methods). Number of stimulus events (from left to right) are: 426, 161, 472, 99, 441, 102, 787, 203. Error bars represent 95% CI. Kolmogorov–Smirnov test was performed to test statistical significance. Significance levels are chosen to account for Bonferroni multiple hypothesis correction. D) Distribution in change in velocities reported in (C) are shown as a probability density.

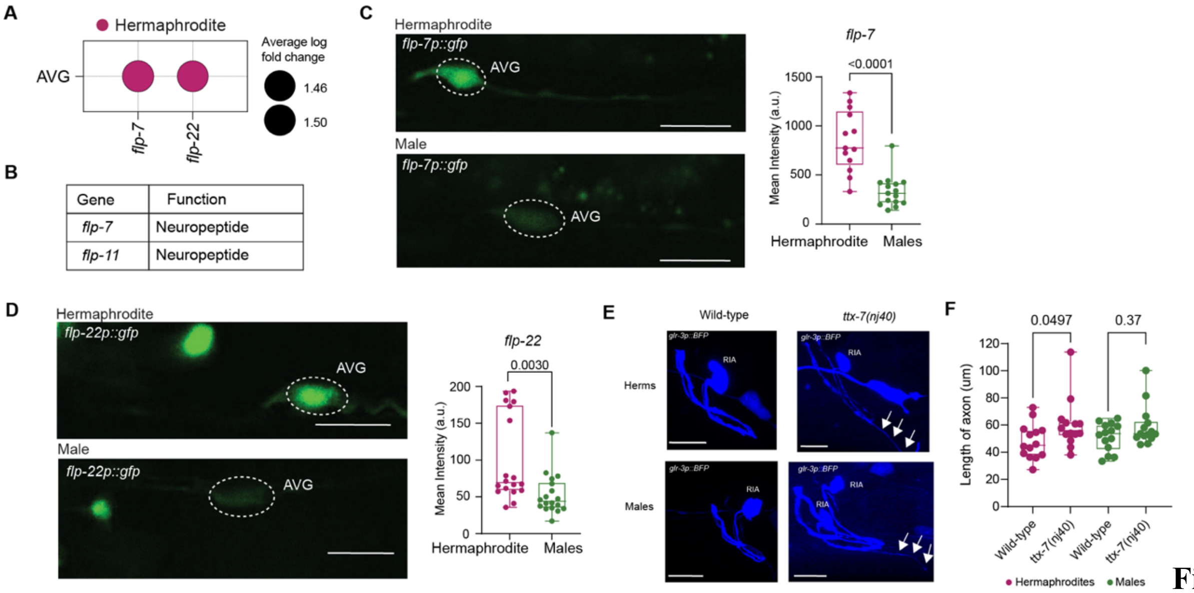


**Fig. S9. Neuropeptides expression in AVG and RIA axons in *ttx-7* mutants**

Bubble plot representation of *flp-7* and *flp-22* genes expression in AVG clusters of the two sexes. Bubble size represents average log fold change, bubble color pink represents hermaphrodites). (B) Table listing the established functions of *flp-7* and *flp-22* genes. (C) Representative confocal micrographs showing *flp-7p::GFP* expression in AVG neurons (white dotted circle) (left). Quantification of *flp-7p::GFP* (right). n = 13 hermaphrodites and n= 15 male worms. (D) Representative confocal micrographs showing *flp-22p::GFP* expression in AVG neurons (white dotted circle) (left). Quantification of *flp-22p::GFP* (right). n = 18 animals per group. a.u., arbitrary units. Scale bars represent 10 µm. (E) Representative confocal micrographs showing *glr-3p::BFP* expression labeling RIA neurons in Wild type (left) and in *ttx-7(nj40)* mutant animals (right) of hermaphrodites (top panels) and males (bottom panels). White arrows represent RIA axon elongation. Scale bars represent 10 µm. (F) Quantification of RIA axon length (um) in hermaphrodites (magenta) and in males (green) in Wild-type and in *ttx-7(nj40)* mutant animals. n = 14 animals per group. In the box-and-whiskers graph, the center line in the box denotes the median, while the box contains the 25th to 75th percentiles of the dataset, whiskers define the minimum and maximum value with dots showing all points. We performed a two-sided Mann-Whitney test for each comparison.

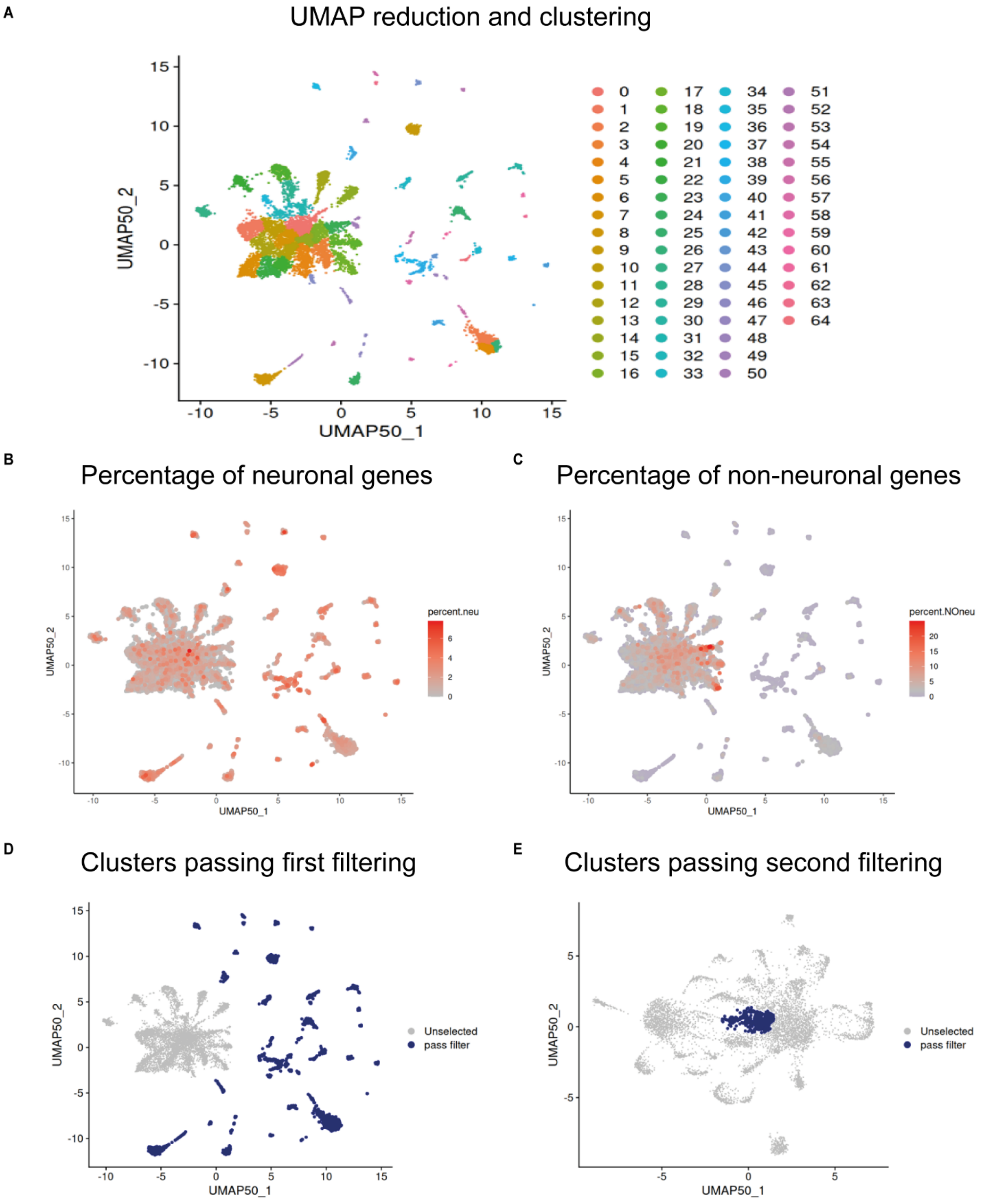


**Fig. S10. Example of neuronal cell filtering applied to a male sample batch**

(A) UMAP reduction using 50 PCAs and clustering with Louvain algorithm using Seurat. (B) Percentage of neuronal and non-neuronal (C) genes shown over UMAP space. (D) First filtering showing clusters passing the threshold of 0.5% Neuronal markers and 1.5% non-Neuronal markers in blue (4012 cells passing the filter) and in gray cells that will be used for a second round of filtering. (E) Second round of filtering were only 365 cells were retained as neurons. In this example a total of 4377 from 11152 cells were kept for the merge of the experimental batches.

**Supplementary Tables S1-S10**

Table S1: Matrix of neuronal and non-neuronal markers used to assign tissue and cell type.

Table S2: All single-cell clusters obtained for both sexes.

Table S3: List of differentially expressed genes between hermaphrodite and male.

Table S4: 10x Genomics 3' Single-cell Experiment Details.

Table S5: List of the 62 sex-shared neuronal clusters that passed all the filters.

Table S6: Binary expression patterns for all clusters based on threshold 2 (0.04).

Table S7: Mean expression and fraction values for annotated clusters of hermaphrodites.

Table S8: Mean expression and fraction values for annotated clusters of males.

Table S9: Number of synapses calculated for Figure 4C-D and Figure 7.

Table S10: Strains, reagents and resources used in this study.
